# Abnormal morphology biases haematocrit distribution in tumour vasculature and contributes to heterogeneity in tissue oxygenation

**DOI:** 10.1101/640060

**Authors:** Miguel O. Bernabeu, Jakub Köry, James A. Grogan, Bostjan Markelc, Albert Beardo, Mayeul d’Avezac, Romain Enjalbert, Jakob Kaeppler, Nicholas Daly, James Hetherington, Timm Krüger, Philip K. Maini, Joe M. Pitt-Francis, Ruth J. Muschel, Tomás Alarcón, Helen M. Byrne

## Abstract

Oxygen heterogeneity in solid tumours is recognised as a limiting factor for therapeutic efficacy. This heterogeneity arises from the abnormal vascular structure of the tumour, but the precise mechanisms linking abnormal structure and compromised oxygen transport are only partially understood. In this paper, we investigate the role that RBC transport plays in establishing oxygen heterogeneity in tumour tissue. We focus on heterogeneity driven by network effects, which are challenging to observe experimentally due to the reduced fields of view typically considered. Motivated by our findings of abnormal vascular patterns linked to deviations from current RBC transport theory, we calculate average vessel lengths 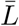 and diameters 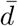 from tumour allografts of three cancer cell lines and observe a substantial reduction in the ratio 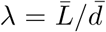 compared to physiological conditions. Mathematical modelling reveals that small values of the ratio *λ* (*i.e. λ* < 6) can bias haematocrit distribution in tumour vascular networks and drive heterogeneous oxygenation of tumour tissue. Finally, we show an increase in the value of *λ* in tumour vascular networks following treatment with the anti-angiogenic cancer agent DC101. Based on our findings, we propose *λ* as an effective way of monitoring the efficacy of antiangiogenic agents and as a proxy measure of perfusion and oxygenation in tumour tissue undergoing anti-angiogenic treatment.

**Significance statement:** Oxygen heterogeneity in solid tumours is recognised as a limiting factor for therapeutic efficacy. This heterogeneity arises from the abnormal tumour vascular structure. We investigate the role that anomalies in RBC transport play in establishing oxygen heterogeneity in tumour tissue. We introduce a metric to characterise tumour vasculature (mean vessel length-to-diameter ratio, *λ*) and demonstrate how it predicts tissue oxygen heterogeneity. We also report an increase in *λ* following treatment with the antiangiogenic agent DC101. Together, we propose *λ* as an effective way of monitoring the action of anti-angiogenic agents and a proxy measure of oxygen heterogeneity in tumour tissue. Unravelling the causal relationship between tumour vascular structure and tissue oxygenation will pave the way for new personalised therapeutic approaches.

## 1 Introduction

Tissue oxygenation plays a crucial role in the growth and response to treatment of cancer. Indeed, well-oxygenated tumour regions respond to radiotherapy better than hypoxic, or oxygen-deficient, regions by a factor of up to three [1, 2]. Further, the increased rates of proteomic and genomic modifications and clonal selection associated with anoxia (*i.e.* total oxygen depletion) endow tumours with more aggressive and metastatic phenotypes [3, 4]. Oxygen heterogeneity in solid tumours is commonly attributed to their abnormal vasculature [5, 6]. This link is arguably multifactorial, including nonuniform vessel distribution, inefficient vessel organisation (*e.g.* functional shunting), flow fluctuations, and variations in red blood cell flux (see [7] for a review). However, the dynamics of abnormal oxygen transport at whole vascular network level have not been addressed.

Oxygen is transported through the vasculature by binding to haemoglobin in red blood cells (RBCs) [8]. Haematocrit, or the volume fraction of RBCs in whole blood, does not distribute uniformly throughout the vasculature [9, 10]. At a bifurcation with one afferent and two efferent branches, it is typically assumed that the efferent branch with the highest flow rate will have the highest discharge haematocrit [10, 11] due to, among other features, plasma-skimming caused by the presence of an RBC-depleted layer or cell free layer (CFL) [12]. In addition, confinement effects can lead to further reduction of the tube haematocrit compared to the discharge value. In this study, we will refer to discharge haematocrit only. How abnormal tumour vascularisation impacts haematocrit splitting (HS) at bifurcations and the potential network effects arising have not been addressed.

In this paper, we investigate the role that anomalies in RBC transport play on establishing oxygen heterogeneity in tumour tissue. We focus on heterogeneity driven by network effects, which are difficult to resolve experimentally due to the reduced fields of view typically considered. Live imaging of tumour allografts of three cancer cell lines reveal abnormal morphological vascular patterns such as short inter-bifurcation distances and complex topological configurations. Furthermore, detailed numerical simulations describing the transport of RBCs predict deviations from current HS theory in these scenarios. When we extract average vessel lengths 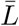 and diameters 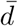 from tumour allografts of three cancer cell lines we observe a substantial reduction in the ratio 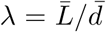, an accepted parameter governing CFL dynamics, compared to physiological conditions. In addition, our RBC simulations reveal: a) asymmetric CFL width disruption following a bifurcation, and b) that the average measured *λ* value in tumour allografts is too small for the CFL to recover full symmetry between consecutive branching points, leading to uneven haematocrit split in the downstream branching point.

Based on the RBC simulations, we propose a new haematocrit splitting rule that accounts for CFL disruption due to pathologically small *λ* values. We integrate this rule into existing models of tumour blood flow and oxygen transport [13] and observe a haematocrit memory effect leading to haemoconcentration/haemodilution in densely branched vessel networks. As a consequence, the predicted tissue oxygenation is highly heterogeneous and differs markedly from predictions generated using rules for HS under physiological conditions (*e.g.* [14, 15, 16, 5, 17]). These findings provide a mechanistic explanation for previous reports of haemoconcentration/haemodilution in tumour vasculature [18], plasma flow [19], and well-perfused vessels that are hypoxic [20]. Furthermore, previous work has demonstrated that fluctuations in red blood cell flux in the tumour microvasculature are associated with instabilities in oxygen concentration in the extravascular space [21, 22]. However, the mechanism underlying these fluctuations remains unclear. Our findings of haematocrit network effects support the hypothesis that vascular structural instability (associated with vascular remodelling or angiogenesis) can induce long-range changes in red blood cell flux throughout the network.

Finally, we demonstrate the preclinical relevance of our findings by showing an increase in the average *λ* value of tumour vascular networks following treatment with the DC101 anti-angiogenic cancer agent. Based on our results, we propose *λ* as an effective way of monitoring the action of anti-angiogenic agents and as a proxy measure of oxygen heterogeneity in tumour tissue undergoing anti-angiogenic treatment. Future experimental studies should confirm this finding and elucidate its relative importance compared to well-characterised mechanisms of normalisation (*e.g.* permeability reduction, vessel decompression [23]).

## 2 Results

### 2.1 Average distance between vessel branch points is shorter in solid tumours than in healthy tissue

We implemented a protocol for *in-vivo* imaging of tumour vasculature [24] and exploited our recently published methods for vessel segmentation [25, 26] and three-dimensional (3D) vascular network reconstruction to characterise the morphology of tumour vasculature. Most analyses of tumour vasculature are based on histological sections that give only partial information about the (3D) structure of the entire network. Vascular casts (see [27] for a review) and optical clearance techniques [28] have been used successfully to image the complete tumour vasculature but they provide information at only one time point. Further, information about vessel perfusion and/or the physiological pressures that drive blood flow are not available. By applying intravital video microscopy to mice in which all the endothelium is flourescent, it was possible to image the entire tumour vasculature: individual vessels could be detected regardless of their level of perfusion (see Methods section for more details). Briefly, tumour allografts of three murine cancer cell lines (*i.e.* MC38, colorectal carcinoma; B16F10, melanoma; and LLC, Lewis lung carcinoma) were implanted in mice, controlled for size, and imaged through an abdominal window chamber using a multi-photon microscope over multiple days. The vascular networks in the 3D image stacks were segmented and the associated network skeletons and vessel diameters computed [25, 26]. Flow direction was determined by tracking the movement of fluorescently labelled RBCs. Upon inspection, the segmented vascular networks presented abnormal morphological vascular patterns such as short inter-bifurcation distances and complex topological configurations such as three-to-two branch mergers (Figure 1(a)–(b)). Motivated by these findings, we exploited recent advances in blood flow simulation methods by our group and others [29, 30, 31, 32] to investigate the link between these abnormal patterns and RBC transport. A computational model of liquid-filled deformable particles (discocytes approximating the shape of an RBC) suspended in an ambient fluid is used to simulate blood flow in a subset of the network, with RBCs inserted at the network inlet and removed at the outlets (see Materials and Methods and Supplementary Information for more details). Our simulations demonstrate that reduced inter-bifurcation distance can lead to an inversion of HS, as predicted by existing theory [11], where the branch with highest flow receives proportionally fewer RBCs and vice versa (Figure 1(c)–(d)). In addition, the same effect is highly attenuated in the complex topology case.

**Figure 1:**
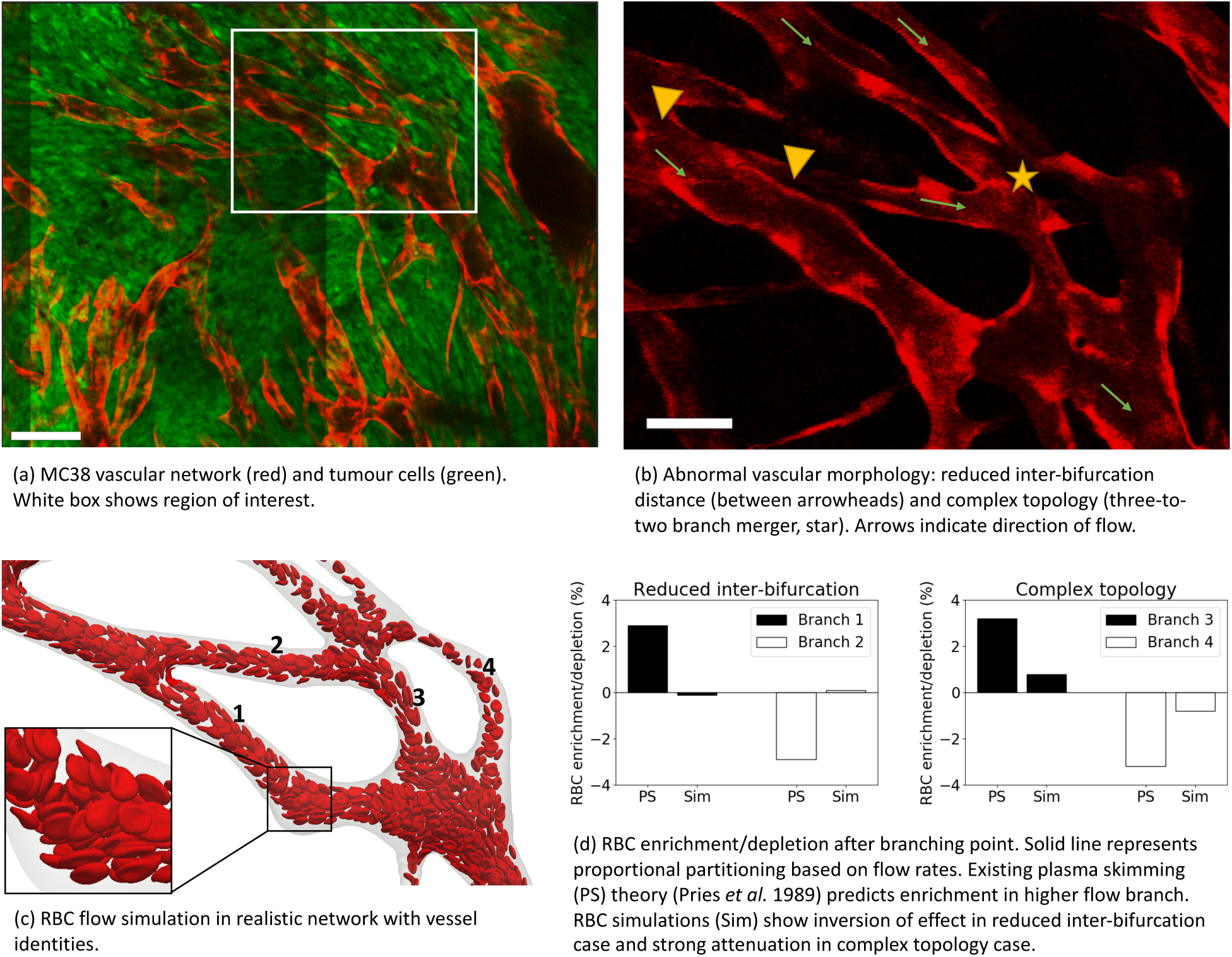
(a) Multiphoton image (0.83 *µ*m × 0.83 *µ*m × 3 *µ*m resolution) of a MC38 tumour vessel network obtained via an abdominal imaging window in mouse. Red — endothelial cells, green — tumour cells. Scale bar: 100 *µ*m. (b) Region of interest showing endothelial cell staining channel alone. Scale bar: 50 *µ*m. (c) RBC flow simulation in computational domain reconstructed from region of interest. (d) RBC flow simulation analysis.

We then asked how prevalent these abnormal vascularisation patterns are throughout the tumour networks imaged. Figure 2(a)–(b) shows the two-dimensional (2D) maximum projection of an example dataset along with a 2D projection of its segmentation and a close-in overlaying segmentation and skeletonisation. Vessel lengths (*L*) and diameters (*d*) in the networks followed a right-skewed distribution resembling a log-normal distribution (Figure 2(c)–(e)). No correlation was found between the variables (Pearson’s *r*^2^ < 0.04 for all samples analysed, Figure 2(c)–(e), Supplementary Tables S4–S5).

**Figure 2:**
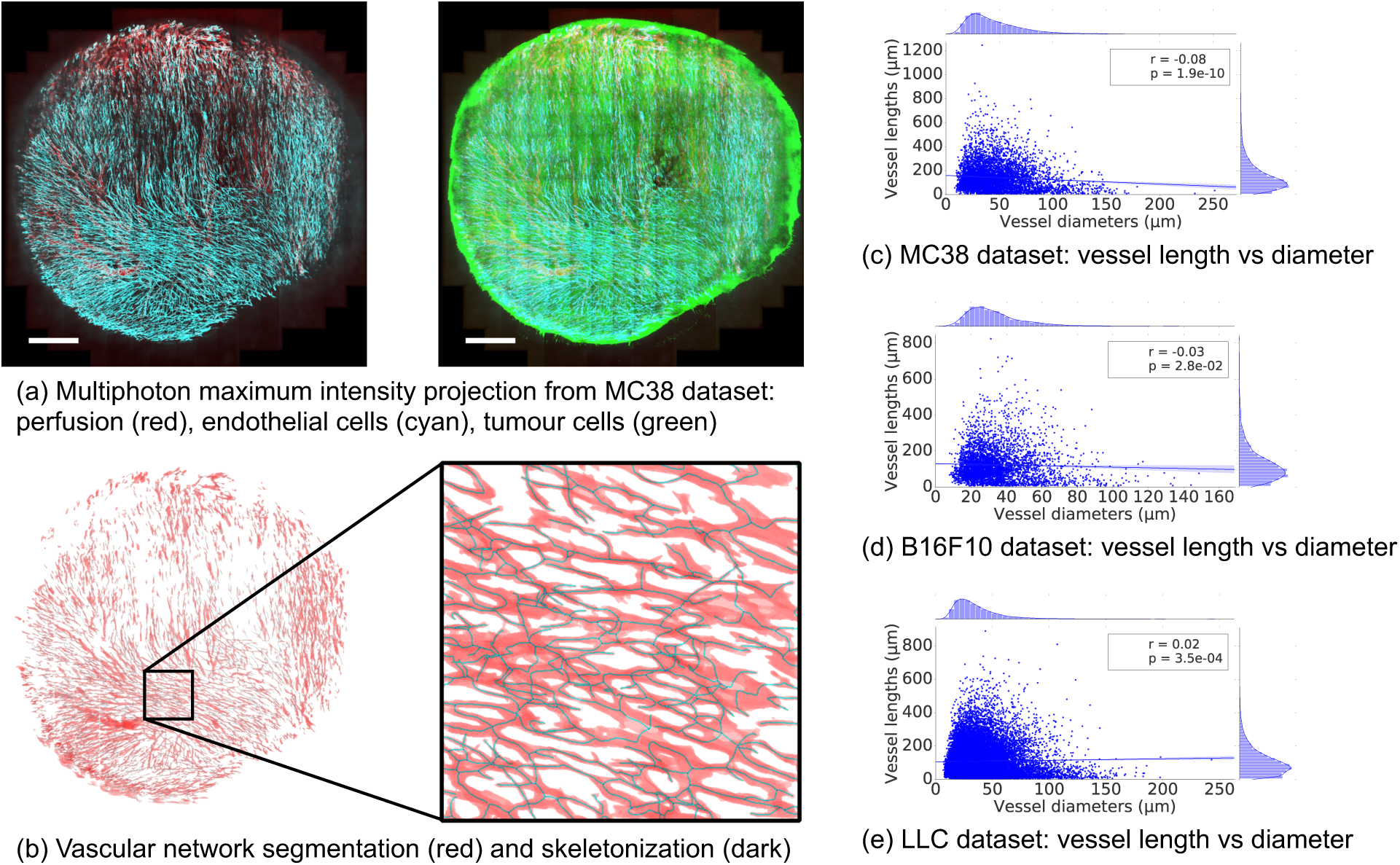
(a) Maximum intensity projection of multiphoton image stack of a tumour vessel network obtained via an abdominal imaging window in mouse. Red — perfusion, Cyan — endothelial cells, Green — GFP tumour cells. Scale bar: 1 mm. (b) The stack is subsequently segmented and skeletonised and distributions of vessel diameters and lengths are calculated. Mouse MC38-5 in Table 1. Scatter plots of vessel lengths versus diameters for different cell lines studied: (c) MC38 (Mouse 3 in Table 1), (d) B16F10 (Mouse 1 in Table 1), (e) LLC (Mouse 1 in Table 1).

Table 1 summarises last-day statistics for all the experiments and averages per cell line. In the example MC38 dataset from Figure 2(a), average vessel length 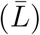 and diameter 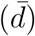 were 143 *µ*m and 45.5 *µ*m, respectively We observe how the group average vessel length is 128.6 *µ*m, 125.9 *µ*m, 108.8 *µ*m for MC38, B16F10, and LLC, respectively. The average diameters are 33 *µ*m, 36.5 *µ*m, 35.7 *µ*m, respectively, which is within the range previously described for tumour vasculature [33]. In addition, the length-to-diameter ratios (*λ*) are 4.0, 3.4, 3.0, respectively, which is substantially smaller than typical *λ* values reported under physiological conditions in a variety of tissues (Supplementary Table S1) and representative of the high branching density encountered in tumour vasculature [34] and in line with the observation in Figure 1(b).

**Table 1:** Mean branch lengths 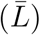, mean vessel diameters 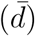, and length-to-diameter ratio 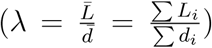 measured over all the imaged vessels in tumour allografts from three murine cancer cell lines. See Supplementary Tables S4–S5 for correlation between variables.

| Cell line | MC38 |  |  |  |  |  |  |
| --- | --- | --- | --- | --- | --- | --- | --- |
| Mouse | 1 | 2 | 3 | 4 | 5 | 6 | Av. |
| $\bar{L}, \mu\text{m}$ | 123.9 | 129.9 | 143 | 112.4 | 132.1 | 130.0 | 128.6 |
| $\bar{d}, \mu\text{m}$ | 29.3 | 33.0 | 45.5 | 23.9 | 28.9 | 37.5 | 33.0 |
| $\lambda = \bar{L}/\bar{d}$ | 4.2 | 3.9 | 3.1 | 4.7 | 4.6 | 3.4 | 4.0 |

| Cell line | B16F10 |  |  |  | LLC |
| --- | --- | --- | --- | --- | --- |
| Mouse | 1 | 2 | 3 | Av. | 1 |
| $\bar{L}, \mu\text{m}$ | 123.2 | 123.5 | 131.0 | 125.9 | 108.8 |
| $\bar{d}, \mu\text{m}$ | 33.9 | 34.1 | 41.6 | 36.5 | 35.7 |
| $\lambda = \bar{L}/\bar{d}$ | 3.6 | 3.6 | 3.1 | 3.4 | 3.0 |

### 2.2 Plasma skimming in tumour-like vasculature is biased by history effects arising from cell free layer dynamics

Our finding of reduced inter branching point distance in tumour tissue motivated us to investigate a potential causal relationship between the reduction in *L* and *λ* and the profoundly abnormal tumour haemodynamics and mass transport patterns described in the literature [35]. Of particular interest is establishing whether haemorheological phenomena may contribute to tumour heterogeneity and hypoxia.

The presence of an RBC-depleted region adjacent to the vessel walls (*i.e.* the cell free layer (CFL)) is a key contributor to plasma skimming (PS) [10, 11, 12]. Previous studies have shown CFL disruption after microvascular bifurcations and found that the length required for CFL recovery, *l*_*r*_, is in the region of 10 vessel diameters (*d*) for *d* < 40*µ*m [11], 8–15*d* for *d* ∈ [20, 24]*µ*m [36], and 25*d* for *d* ∈ [10, 100]*µ*m [37]. These values are substantially higher than the average *λ* values given in Table 1, *λ* < *l*_*r*_, and therefore we expect that, on average, CFL symmetry will not recover between the branching points in the networks under study.

Motivated by these findings, we applied our RBC simulation approach to investigate the link between CFL dynamics and PS in a tumour-inspired 3D microvascular network. Our intention is to understand the extent to which CFL disruption effects arising at a bifurcation affect haematocrit splitting in downstream bifurcations for small interbifurcation distances relevant to tumour vasculature (see Methods section for further details). Briefly, we define a set of networks of 3D cylindrical channels of constant radius, consisting of one main channel with an inlet and an outlet and two side branches, which define two consecutive bifurcations (Figure 3). We consider inter-bifurcation distances of four and 25 channel diameters based on our tumour vascular network analysis and the largest of the CFL recovery distances reviewed earlier. We position the two side branches on the same side of the main channel or on opposite sides. Flow rates at the network inlet and outlets are configured such that at each bifurcation flow is split evenly. We perform blood flow simulations (3 runs in each network, with random perturbations in the RBC insertion procedure) and, after the initial transient time required to fully populate the network with RBCs, we quantify discharge haematocrit by an RBC-counting procedure.

**Figure 3:**
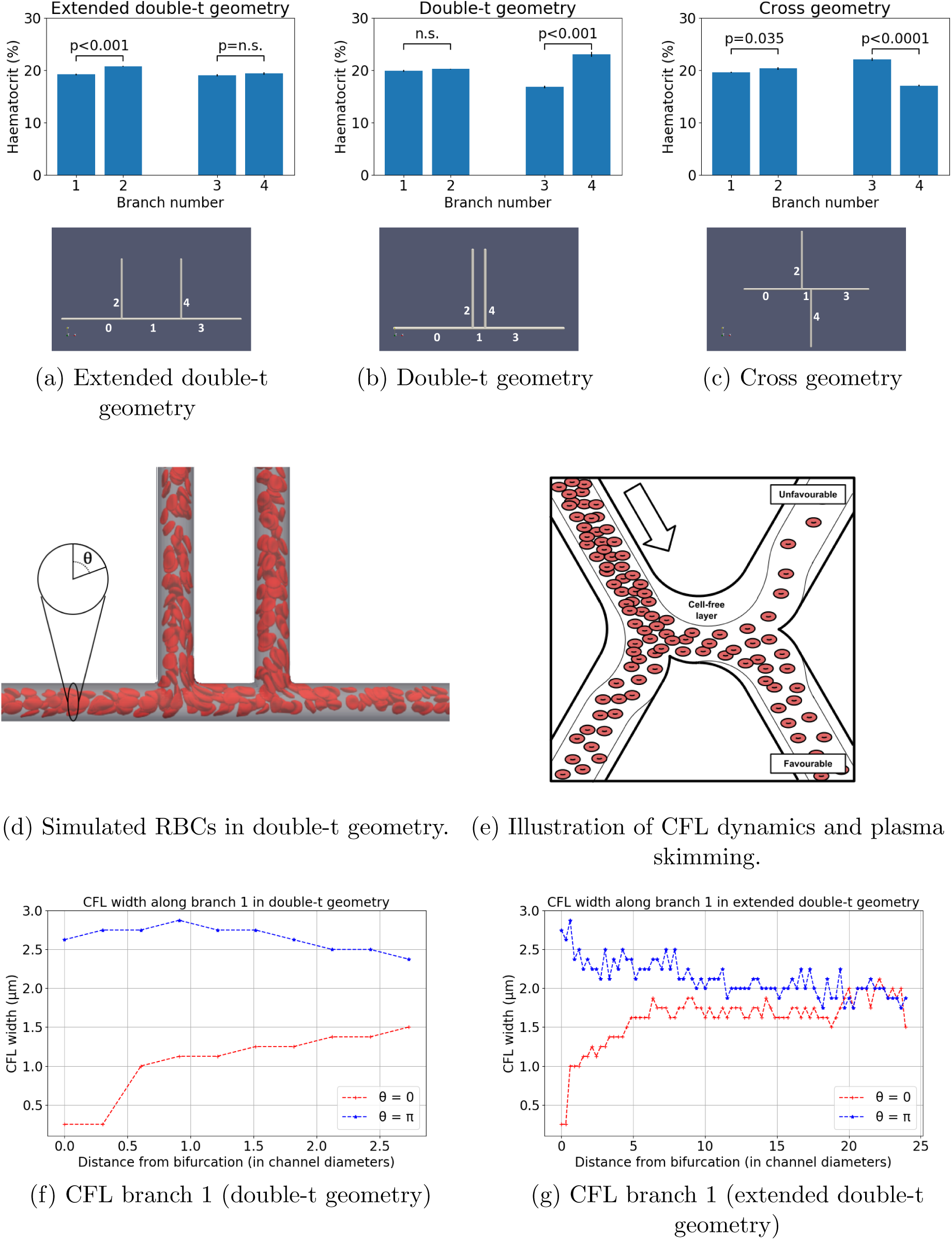
Discharge haematocrit at different geometry branches: (a) extended double-t geometry, (b) double-t geometry, (c) cross geometry. Example simulation in the double-t geometry: (d) vessel network is rendered semi-transparent in grey, RBC membranes are rendered in red suspended in transparent blood plasma. (e) Schematic describing the impact of CFL dynamics on haematocrit split. CFL width in opposite sides of channel 1: (f) double-t geometry, (g) extended double-t geometry.

Figure 3a–3b and Table 2 show how haematocrit split is close to even at bifurcation 1 for all geometries studied, as would be predicted by existing theoretical models of HS. However, different degrees of haematocrit splitting occur at bifurcation 2. In the doublet geometry, we observe haemodilution in branch 3 and haemoconcentration in branch 4 (16.8% vs 23%, *p* < 0.001, Figure 3b), which we will refer to as the unfavourable and favourable branches. These effects are not statistically significant in the same branches in the extended double-t geometry (19.1% vs 19.4%, *p* = 0.3, Figure 3a). The haemoconcentration/haemodilution effect is present in the cross geometry but the branches experiencing it are interchanged (22.1% vs 17.1%, *p* < 0.001, Figure 3c). In contrast with these results, existing HS theoretical models would predict even haematocrit splitting at bifurcation 2, regardless of the inter-bifurcation distance, due to the prescribed symmetrical flow and geometry conditions.

**Table 2:** Discharge haematocrit calculated at the different branches of each bifurcation for the extended double-t (EDT) geometry, double-t (DT) geometry, and cross (X) geometry. Values are given as mean (standard error) over an ensemble of three simulations with random perturbations in the RBC insertion procedure while the haematocrit at the inlet is held constant.

|  | Bifurcation 1 |  |  |
| --- | --- | --- | --- |
|  | Branch 0 | Branch 1 | Branch 2 |
| EDT | 20.06(0.03) | 19.23(0.14) | 20.80(0.10) |
| DT | 20.08(0.06) | 19.96(0.19) | 20.26(0.05) |
| X | 20.06(0.02) | 19.67(0.09) | 20.35(0.20) |

|  | Bifurcation 2 |  |  |
| --- | --- | --- | --- |
|  | Branch 1 | Branch 3 | Branch 4 |
| EDT | 19.23(0.14) | 19.05(0.22) | 19.4(0.23) |
| DT | 19.96(0.19) | 16.83(0.24) | 23.04(0.46) |
| X | 19.67(0.09) | 22.12(0.26) | 17.09(0.13) |

On closer inspection, the dynamics of the CFL show how, after bifurcation 1, CFL width is initially negligible and rapidly increases on the side of channel 1 leading to the favourable branch (*θ* = 0, Figure 3f). Conversely, CFL width increases after the bifurcation and follows a downward trend in the opposite side (*θ* = *π*, Figure 3f). An inter-bifurcation distance of four diameters is too short for the CFL width to equalise on both sides (Figures 3f). In contrast, CFL width has time to become symmetric on both sides for an inter-bifurcation distance of 25 diameters (Figure 3g).

Taken together, these results show how CFL asymmetry can cause uneven haematocrit split in bifurcation 2 (Figure 3e) irrespective of branching side, *i.e.* cross *vs* double-t geometry. Our results are consistent with the findings by Pries *et al.*, describing how asymmetry of the haematocrit profile in the feeding vessel of a bifurcation has a significant influence on RBC distribution in the child vessels [11]. In addition, we provide quantitative evidence of how CFL asymmetry may be the main contributing factor.

Interestingly, we observe small but statistically significant asymmetries in the haematocrit split in bifurcation 1 in the extended double-t geometry (19.2% vs 20.8%, *p* < 0.001, Figure 3a) and cross geometry (19.7% vs 20.4%, *p* = 0.035, Figure 3c), which consistently favour the side branch. We attribute this secondary effect to an asymmetrical streamline split in the chosen geometry as investigated in [38].

We note that the effects described above depend on the angle between the planes containing the two consecutive bifurcations. Our data suggest that for angles of 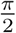 radian the asymmetric haematocrit split effects will not be observed since the CFL width at 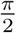 and 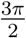 remains mostly symmetric (Supplementary Figure S2).

### 2.3 Haematocrit history effects lead to highly heterogeneous oxygen distributions in solid tumours

Existing theoretical models of haematocrit splitting (HS) [11, 39, 40, 41] do not capture the haemoconcentration/haemodilution effects in realistic and idealised vascular networks presented in Figures 1 and 3. We hypothesise that this is because they neglect CFL disruption at bifurcations and its impact on subsequent bifurcations. We propose a new HS model which accounts for CFL dynamics and show that it predicts history effects in dense networks (see Materials and Methods section for details and Supplementary Information for a description of its validation). The new model is significantly less computationally intensive to solve than the RBC simulations (see Materials and Methods for details).

We use Murray’s law [42] and our experimentally measured values of *λ* to design a synthetic vessel network comprising consecutive double-t/cross bifurcations (see Supplementary Figure S4a and Materials and Methods for details). Most notably, at all bifurcations the flow split and the radii of the child vessels are equal, a scenario where existing HS models would predict homogeneous haematocrit throughout the network. We simulate network blood flow using a Poiseuille flow approximation with a HS model originally proposed by Pries *et al.* [11, 39] (without memory effects) and our new model (accounting for memory effects). As for the RBC simulations, differences in haematocrit between child branches emerge after two bifurcations (Supplementary Figure S4c), and are amplified with increasing vessel generation number (Supplementary Figure S4d).

Our model predicts the emergence of a compensatory mechanism in child branches. Increased flow resistance in the branch experiencing haemoconcentration leads to partial re-routing of flow in the other branch (Supplementary Figure S4b). This, in turn, attenuates the haemoconcentration/haemodilution effects previously described due to HS dependence on flow ratios.

We now consider how this memory effect in the haematocrit distribution may affect oxygen distribution in the tissue being perfused by the synthetic network. Following [13] (see Materials and Methods for a description of the coupled model), the haematocrit distribution in the network acts as a distributed source term in a reaction-diffusion equation for tissue oxygen. We define sink terms so that oxygen is consumed at a constant rate everywhere within the tissue. The equation is solved numerically and oxygen distributions generated using two HS models (with and without memory effects, Supplementary Figures S5a–S5b) are compared for a range of *λ* values. We focus on the central portion of the network (Figures 4a–4b) where the tissue is densely vascularised. The results presented in Figure 4c and Supplementary Figure S6 show that for larger values of *λ* the differences in the oxygen distribution in the tissue for the two HS models are not statistically significant (*e.g.* with *λ* = 10, *p* = 0.14). However, as *λ* decreases, statistically significant differences appear (*e.g.* with *λ* = 4, *p* < 0.001). Without memory effects, the oxygen distributions become narrower as *λ* decreases; with memory effects, the oxygen distributions are increasingly wider for *λ* < 10 (see Supplementary Figure S6c) leading to higher dispersion in the distributions for small *λ*. This is indicative of much more heterogeneous tissue oxygenation.

**Figure 4:**
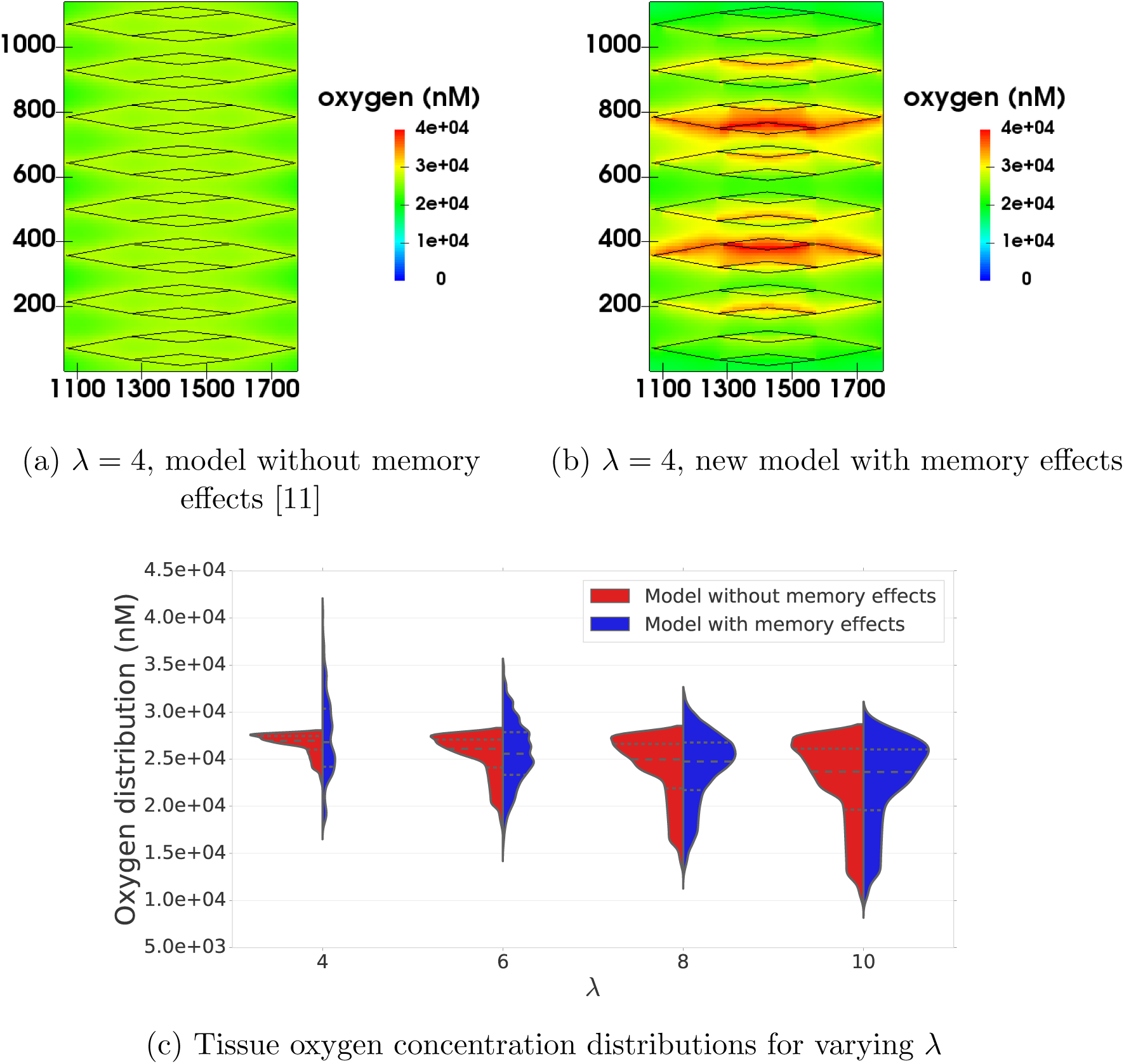
For *λ* = 4.0, the new model with memory effects yields more pronounced oxygen heterogeneity than previous theory [11] predicts (*i.e.* more dispersed oxygen distribution) in (a) and (b) (spatial scales are in microns, vessels shown in black for reference). Violin plots in (c) show oxygen distributions for varying *λ* and the two HS models under consideration. Heterogeneity increases with *λ* for the model without memory effects as expected, but the model with memory effects predicts increased heterogeneity for very low *λ*. The horizontal lines in oxygen distributions in (c) represent 25, 50 and 75% percentiles.

In the current study, we do not consider other sources of heterogeneity such as anisotropic transport and heterogeneous consumption of oxygen or other morphological abnormalities in the vascular networks. We hypothesise that their interaction with the haematocrit history effects reported here will further accentuate tissue oxygen heterogeneity.

### 2.4 Vascular normalisation therapies increase *λ* ratio in tumours

Our findings of reduced *λ* ratio in tumour vasculature and associated predictions of increased oxygen heterogeneity led us to investigate whether existing vascular normalisation therapies modulate this parameter. Previous reports (Supplementary Table S2) have extensively demonstrated in multiple animal models that anti-angiogenic treatment leads to reduction in tumour vessel diameters. In those studies that analyse vessel length and diameter post-treatment, vessel length either remains unchanged or decreases to a lesser extent than vessel diameter. These findings indicate an increase in *λ* ratio post-treatment. Furthermore, Kamoun *et al.* also reported a reduction in tumour haemoconcentration post-treatement [43], which suggests an *in-vivo* link between an increase in *λ*, haematocrit normalisation and oxygen transport homogenisation.

We validated these results in our animal model by calculating the *λ* ratio following DC101 treatment (see Materials and Methods for details). Our results indicate that in the first two days post-treatment *λ* increases significantly and then starts to decrease matching the control trend (Figure 5a, Supplementary Table S3). This change is explained by a linear increase in vessel length immediately after treatment (absent in the control group), which is compensated after two days by an increase at a higher rate in vessel diameter (comparable to the control group) (Supplementary Figure S7), and associated with a decrease in bifurcation density as would be expected from impaired angiogenesis and vessel pruning (Figure 5b, Supplementary Figure S7). Our results also demonstrate that, for vessels with diameters less than 50 *µm*, the fraction of perfused vessels increases after DC101 treatment, reaching a maximum of 63% relative increase compared to control after two days, before decreasing to match the controls after three days (Figure 5c). We note that the proportion of vessels with diameter less than 50 *µm* is identical between groups, regardless of perfusion (Supplementary Figure S7). Interestingly, the time window in which perfusion peaks coincides with the largest, post-treatment increase in *λ*. However, the structural changes persist beyond this window, which would suggest that improvements in oxygen transport homogeneity can be independent of perfusion.

**Figure 5:**
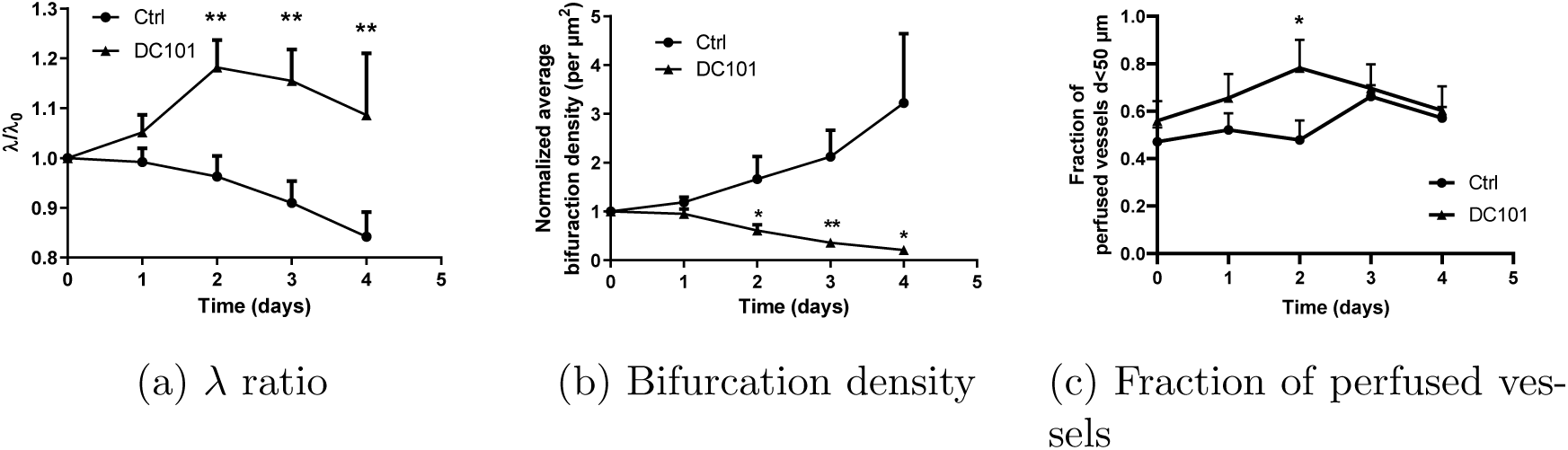
Vascular phenotypes in MC38 tumours over time following DC101 treatment compared with control (n=5). DC101 *λ* ratio raw data is given in Supplementary Table S3. * *p* < 0.05, ** *p* < 0.01.

## 3 Discussion

Hypoxia compromises the response of many tumours to treatments such as radiotherapy, chemotherapy, and immunotherapy. Dominant causative factors for hypoxia associated with the structure and function of the tumour vasculature include inefficient vessel organisation and inadequate flow regulation. Motivated by morphological analyses of vascular networks from different tumour types and detailed computer simulations of RBC transport through realistic and synthetic networks, we have proposed a new, rheological mechanism for tumour hypoxia.

We analysed vascular networks from murine MC38, B16F10, and LLC tumour allografts. Upon inspection, the segmented vascular networks presented abnormal morphological patterns such as reduced inter-bifurcation distance and branching points with topological configurations different from the typical diverging/converging bifurcations. Detailed computer simulations of RBC transport uncovered a link between these morphological anomalies and strong deviations from the current HS theory, including inversion of HS (a phenomenon that had only been previously observed *in vitro* [44, 45]). For each vessel segment within each network, we calculated the metric *λ*, which is the ratio of its length and diameter, and has been shown to control CFL dynamics. Average *λ* values for the three tumour cell lines were similar in magnitude 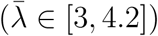 and several fold smaller than values from a range of healthy tissues 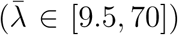. Our RBC simulations confirmed previous reports of transient alterations in the CFL downstream of network bifurcations (*e.g.* asymmetries in the cross-sectional haematocrit profile following a bifurcation [46, 47] and the temporal dynamics governing its axisymmetry recovery [37]). Further, for the *λ* values measured in our tumours and the capillary number considered in our simulations, the CFL did not become symmetric between consecutive branching points. This bias is amplified across branching points and drives haemoconcentration/haemodilution at the network level. Based on these findings, we developed a new rule for haematocrit splitting at vessel bifurcations that accounts for CFL disruption due to abnormally short vessel segments and validated it against our fully resolved RBC simulations. We then used our existing oxygen transport model [13] to demonstrate that this haematocrit memory effect can generate heterogeneous oxygen distributions in tissues perfused by highly branched vascular networks and that the network metric *λ* controls the extent of this heterogeneity. Finally, we reported an increase in the average *λ* value of tumour vascular networks following treatment with the DC101 anti-angiogenic cancer agent.

The implications of our findings are multiple. We have introduced a simple metric to characterise tumour vasculature based on the mean length-to-diameter ratio of vessel segments (*λ*), and demonstrated how it controls oxygen heterogeneity in a synthetic, densely vascularised, tissue model. Our findings, of structurally induced haemodilution in vascular networks with low *λ* values, provide a mechanistic explanation for experimental observations of haemodilution in tumour vascular networks [18], plasma flow [19], and the existence of well-perfused vessels that are hypoxic [20]. Furthermore, these findings support the hypothesis that vascular structural instability (associated with vascular remodelling or angiogenesis) leads to the fluctuations in red blood cell flux previously linked with cycling hypoxia [21, 22]. We conclude that vessel perfusion is a poor surrogate for oxygenation in tissue perfused by vascular networks with low *λ* values. Further, predictions of tissue oxygenation based on diffusion-dominated oxygen transport (*e.g.* [14, 15, 16, 5, 17]) may be inaccurate if they neglect heterogeneity in the haematocrit distribution of the vessel network.

Finally, anti-angiogenic drugs have been shown to generate transient periods of heightened homogeneous tissue oxygenation, due to improved restructuring and reduced permeability of tumour vessels [48]. This phenomenon, termed ‘vascular normalisation’ [6], can correct the deficient transport capabilities of tumour vasculature, homogenise drug and oxygen coverage, and, thereby, improve radiotherapy and chemotherapy effectiveness [2]. Our results demonstrate an increase in the average *λ* value of tumour vascular networks post treatment with anti-angiogenic drugs and predict that this would lead to a less heterogeneous haematocrit distribution and more uniform intratumoural oxygenation potentially independent of perfusion. Based on our findings, we propose *λ* as an effective way of monitoring the action of anti-angiogenic agents. Further experimental work, quantifying haematocrit network dynamics before and after anti-angiogenic treatment, is needed to test this prediction and elucidate its importance in comparison with established mechanisms of normalisation (*e.g.* permeability reduction, vessel decompression [23]). If confirmed, this finding would provide a theoretical foundation for the development of therapeutic approaches for the normalisation of tumour oxygenation involving the administration of agents that target *λ* and, therefore, homogenise haematocrit and tissue oxygenation.

## 4 Materials and Methods

### 4.1 Tumour allograft model and abdominal imaging window protocol

The abdominal window chamber was surgically implanted in transgenic mice on C57Bl/6 background that had expression of red fluorescent protein tdTomato only in endothelial cells. The murine colon adenocarcinoma - MC38, murine melanoma - B16F10, and murine Lewis Lung Carcinoma – LLC tumours with expression of green fluorescent protein (GFP) in the cytoplasm were induced by injecting 5 *µ*l of dense cell suspension in a 50/50 mixture of saline and matrigel (Corning, NY, USA). For DC101 treatment, mice bearing MC38 tumours were treated with anti-mouse VEGFR2 antibody (clone DC101, 500 *µ*g/dose, 27 mg/kg, BioXCell) injected intraperitoneally on the first and fourth day of imaging. Prior to imaging we intravenously injected 100 *µ*l of Qtracker 705 Vascular Labels (Thermo Fisher Scientific, MA, USA) which is a blood-pool based labelling agent, thus allowing us to determine whether vessels were perfused or not.

The direction of blood flow in the vessels was determined by tracking the movement of fluorescently labelled RBCs. RBCs were isolated from blood taken from an anesthetised donor mouse via cardiac puncture. RBCs were washed twice in phosphate-buffered saline (PBS) and then labelled with DiD Cell-Labeling Solution (Thermo Fisher). The labelled RBCs were then washed twice again, resuspended in 1 mL of PBS and then 100 *µ*L of the resuspended labelled RBCs was injected via a tail vein prior to imaging. Isoflurane inhalation anesthesia was used throughout the imaging, mice were kept on a heated stage and in a heated chamber and their breathing rate was monitored.

Tumour images were acquired with Zeiss LSM 880 microscope (Carl Zeiss AG), connected to a Mai-Tai tunable laser (Newport Spectra Physics). We used an excitation wavelength of 940 nm and the emitted light was collected with Gallium Arsenide Phosphide (GaAsP) detectors through a 524–546 nm bandpass filter for GFP and a 562.5– 587.5 nm bandpass filter for tdTomato and with a multi-alkali PMT detector through a 670–760 nm bandpass filter for Qtracker 705 and DiD. A 20x water immersion objective with NA of 1.0 was used to acquire a Zstacks-TileScan with dimensions of 512 *×* 512 pixels in *x* and *y*, and approximately 70 planes in *z*. Voxel size was 3–5 *µ*m in the *z* direction and 0.83 *µ*m *×* 0.83 *µ*m in the *x*-*y* plane. Each tumour was covered by approximately 100 tiles. The morphological characteristics of tumour vasculature were obtained from the acquired images as previously described [25, 26]. All animal studies were performed in accordance with the Animals Scientific Procedures Act of 1986 (UK) and Committee on the Ethics of Animal Experiments of the University of Oxford.

### 4.2 RBC simulations in realistic and synthetic capillary networks

We created a realistic network of tumour capillaries (Supplementary Figure S1) based on the images in Figure 1 and a protocol previously described [49, 50]. Flow direction at the boundaries was assigned in agreement with experimental observations of flourescent RBC tracking. The inlets in the domain were connected together to facilitate definition of flow and RBC boundary conditions (and similarly with the outlets).

For our synthetic network simulations, we define a set of networks of cylindrical channels of diameter *d*. An inlet channel of length 25*d* (channel 0) bifurcates into two channels of length *δ* and 25*d* at *π* and *π/*2 radians clockwise, respectively (channels 1 and 2). Channel 1 bifurcates into two channels of length 25*d* at *π* and *α* radians clockwise, respectively (channels 3 and 4). We consider the following configurations (Figure 3): double-t geometry (*δ* = 4*d, α* = *π/*2), cross geometry (*δ* = 4*d, α* = 3*π/*2), and extended double-t geometry (*δ* = 25*d, α* = *π/*2). Inter-bifurcation distance *δ* is calculated between the centrelines of channels 2 and 4 for consistency with our experimental morphological characterisation.

A model of liquid-filled elastic membranes (discocytes of 8 *µ*m diameter approximating the shape of an RBC) suspended in an ambient fluid is used to simulate blood flow in the networks. We use the fluid structure interaction (FSI) algorithm previously presented and validated by Krüger *et al.* [51], which is based on coupling the lattice Boltzmann method (LBM), finite element method (FEM), and immerse boundary method (IBM). The discocyte membranes are discretised into 500 triangles, which imposes a voxel size of 0.8 *µ*m on the regular grid used in the LBM simulation. The mechanical properties of the membrane are defined to achieve a capillary number (*i.e.* the ratio of viscous fluid stress acting on the membrane and a characteristic elastic membrane stress) of 0.1 in channel 0. The coupled algorithm is implemented in the HemeLB blood flow simulation software [52, 49] (http://ccs.chem.ucl.ac.uk/hemelb). Simulations ran on up to 456 cores of the ARCHER supercomputer taking 11–32 hours. See Supplementary Information for full details.

A constant flow rate of 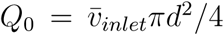 and a procedure for RBC insertion with discharge haematocrit *H*_*inlet*_ is imposed at the network inlet. In the synthetic networks, the outlet flow rates are set to *Q*_2_ = *Q*_0_*/*2 and *Q*_3,4_ = *Q*_0_*/*4 to ensure an equal flow split at each bifurcation. RBCs are removed from the computational domain when they reach the end of any outlet channel. Supplementary Table S6 summarises the key parameters in the model. We performed blood flow simulations (in the synthetic networks 3 runs in each network, with random perturbations in the RBC insertion procedure) and, after the initial transient required to fully populate the network with RBCs, we quantified haematocrit by an RBC-counting procedure.

### 4.3 Hybrid model for tissue oxygen perfusion that accounts for history effects in vascular networks

We first explain how our vascular networks are designed. Then, we describe how blood flow and haematocrit are determined. Next, we introduce the HS models and explain how CFL memory effects are incorporated and the resulting flow problem solved. We conclude by describing how the resulting haematocrit distribution is used to calculate oxygen perfusion in the surrounding tissue. The basic steps of our method are summarised in the flow chart in Supplementary Figure S3.

#### 4.3.1 Synthetic network design

Our networks have one inlet vessel (with imposed blood pressure and haematocrit; we call this generation 0), which splits into two child vessels (generation 1), which then split into two child vessels (generation 2), and so on until a prescribed (finite) number of generations is reached. This defines a sequence of consecutive double-t/cross bifurcations. Thereafter, the vessels converge symmetrically in pairs until a single outlet vessel is obtained (with imposed blood pressure). At every bifurcation, the diameters of the two child vessels are assumed to be equal and determined by appealing to Murray’s law [42]. Using the same vessel diameters in all simulations, we vary vessel lengths, so that for all vessels in the network, the lengths equal the product of *λ* (which is fixed for a given network) and the vessel diameter. We focus on *λ*-values in the range measured in our tumours. We choose a synthetic network so that only *λ*-related effects (and not other morphological network characteristics) contribute to haemoconcentration/haemodilution. In future work, we will investigate the combined effect on tissue oxygenation of the HS model with memory effects and other tumour vascular characteristics.

#### 4.3.2 Blood flow and haematocrit splitting

##### Network flow problem

Tissue oxygenation depends on the haematocrit distribution in the vessel network perfusing the tissue. The haematocrit distribution depends on the blood flow rates. These rates are determined by analogy with Ohm’s law for electric circuits, with the resistance to flow depending on the local haematocrit via the Fahraeus-Lindquist effect (for details, see Supplementary Information and [53]). The flow rates and haematocrit are coupled. We impose conservation of RBCs at all network nodes^1^. A HS rule must then be imposed at all diverging bifurcations.

##### HS model without memory effects

The empirical HS model proposed by Pries *et al.* [11, 54] states that the volume fraction of RBCs entering a particular branch *FQ*_*E*_ depends on the fraction of the total blood flow entering that branch *FQ*_*B*_ as follows:

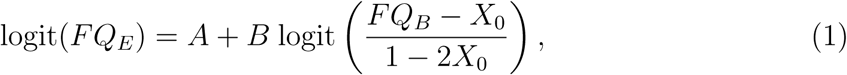

where logit(*x*) = ln (*x/*(1 − *x*)), *B* serves as a fitting parameter for the nonlinear relationship between *FQ*_*E*_ and *FQ*_*B*_, and *A* introduces asymmetry between the child branches (note that for an equal flow split *FQ*_*B*_ = 0.5, *A* =*/* 0 yields uneven splitting of haematocrit). Finally, *X*_0_ is the minimum flow fraction needed for RBCs to enter a particular branch (for lower flow fractions, no RBCs will enter)^2^; the term (1 − 2*X*_0_) reflects the fact that the CFL exists in both child vessels (see Supplementary Figure S8a).

##### HS model with memory effects

We account for the effects of CFL disruption and recovery by modifying the parameters *A* and *X*_0_ (as already observed in [11]). For simplicity, and in the absence of suitable data, we assume that the parameter *B* is the same in both child branches. If *X*_0,*f*_ (*A*_*f*_) and *X*_0,*u*_ (*A*_*u*_) denote the values of *X*_0_ (*A*) in the favourable and unfavourable child branches (see Figure 3e), then our new model of HS can be written as:

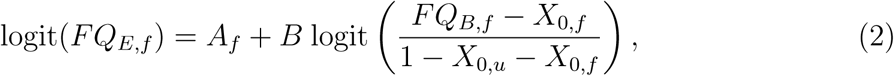

where subscripts _*f*_ and _*u*_ relate to favourable and unfavourable branches, respectively (see Supplementary Figure S8a for a graphical depiction). It is possible to rewrite (2) in terms of the suspension flow rates *Q ≡ Q*_*B*_ and haematocrit levels *H*of the favourable *f*, unfavourable *u*, and parent *P* vessels as (for details see Supplementary Information):

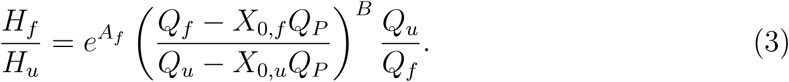

This formulation of our HS model facilitates comparison with other HS models [55, 40, 41]. Functional forms for *A*_*f*_, *X*_0,*f*_ and *X*_0,*u*_ are based on our RBC simulation results and the existing literature (see Supplementary Information). We use an iterative scheme (as in [40]) to determine the flow rates and haematocrit in a given network.

#### 4.3.3 Calculating the tissue oxygen distribution

We embed the vessel network in a rectangular tissue domain. A steady state reaction-diffusion equation models the tissue oxygen distribution, with source terms at vessel network locations proportional to the haematocrit there, and sink terms proportional to the local oxygen concentration modelling oxygen consumption by the tissue. This equation is solved numerically using Microvessel Chaste (see [13] and Supplementary Information for details). In order to highlight the influence of HS on tissue oxygen, we focus on the central 25% of the domain which is well-perfused and ignore the avascular corner regions (see Supplementary Figures S5a and S5b)

### 4.4 Data availability

The source code for the RBC simulations is available at http://ccs.chem.ucl.ac.uk/hemelb. The source code for the oxygen perfusion simulations is available at https://github.com/jmsgrogan/MicrovesselChaste and https://github.com/chaste/chaste.

## Acknowledgements

We acknowledge HZ Ford for the cartoon in Figure 3e, the contributions of the HemeLB development team, and CD Arvanitis and V Vavourakis for helpful comments. Software development was supported by the Engineering and Physical Sciences Research Council (EPSRC) (grant eCSE-001-010). Supercomputing time on the ARCHER UK National Supercomputing Service (http://www.archer.ac.uk) was provided by the “UK Consortium on Mesoscale Engineering Sciences (UKCOMES)” under the EPSRC Grant No. EP/R029598/1. T.K.’s and M.O.B’s contribution have been funded through two Chancellor’s Fellowship at The University of Edinburgh. M.O.B is supported by grants from EPSRC (EP/R029598/1, EP/R021600/1), Fondation Leducq (17 CVD 03), and the European Union’s Horizon 2020 research and innovation programme under grant agreement No 801423. The research leading to these results has received funding from the People Programme (Marie Curie Actions) of the European Unions Seventh Framework Programme (FP7/2007-2013) under REA grant agreement No 625631 (obtained by BM). This work was also supported by Cancer Research UK (CR-UK) grant numbers C5255/A18085 and C5255/A15935, through the CRUK Oxford Centre. This work was supported by the Biotechnology and Biological Sciences Research Council UK Multi-Scale Biology Network, grant number BB/M025888/1. We would like to acknowledge funding from the UK Fluids Network (EPSRC grant number EP/N032861/1) for a Short Research Visit between the Edinburgh and Oxford teams.

## A Supplementary information

The Supplementary Information is organised as follows. First, we provide experimental evidence which supports the findings that vessel lengths and diameters are uncorrelated in tumour environments. Next, we describe the fluid structure interaction (FSI) algorithm used for the red blood cell (RBC) simulations and the method used to calculate the width of the cell free layer (CFL). Next, we present our hybrid model of tissue perfusion and introduce our new haematocrtic splitting (HS) model. Finally, we comment on the higher mean oxygen values predicted by our oxygen perfusion model for small *λ* values.

### A.1 Vessel lengths and diameters in tumour microvasculature are uncorrelated

In Supplementary Tables S4 and S5 we list Pearson’s r-values quantifying the correlation between vessel lengths, *L*, and diameters, *d*,

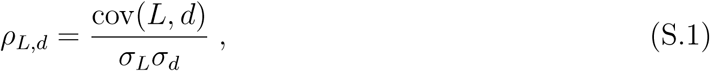

where cov(*i, j*) is the covariance of two variables and *σ*_*i*_ is the standard deviation of variable *i*, for the three tumour cell lines used in our experiments. Results are presented for each mouse and each scan. Day 0 was chosen as the day when the tumour vascular network appeared to be fully formed. This typically occurred approximately 8 days after tumour induction, when the tumour size was approximately 4 mm in diameter. We note also that the duration of the observation period is cell-line specific; some tumours grew faster than others and, as a result, soon started pushing on the window, and in such cases the animal had to be culled as per licence limitations. The Pearson’s r-values are too low to conclude that a correlation exists between *L* and *d* in the tumour vascular networks studied.

### A.2 Red blood cell suspension model

The lattice Boltzmann method (LBM) numerically approximates the solution of the Navier-Stokes equations for a weakly compressible Newtonian fluid discretised on a regular lattice. We employ the D3Q19 lattice, the Bhatnagar–Gross–Krook collision operator extended with the Guo forcing scheme [56], the Bouzidi-Firdaouss-Lallemand (BFL) implementation of the no-slip boundary condition at the walls [57], and the Ladd implementation of the velocity boundary condition for open boundaries [58]. These methods have been extensively used and analysed in the literature (see [59, 60] for a detailed description).

The RBC membrane is modelled as a hyperelastic, isotropic and homogeneous material, following the model described in [29]. The total membrane energy *W* is defined by *W* = *W*^*S*^ + *W*^*B*^ + *W*^*A*^ + *W*^*V*^, where the superscripts denote energy contributions due to strain, bending, area and volume. We employ the surface strain energy density *w*^*S*^ proposed by Skalak *et al.* [61]:

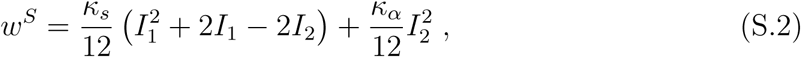

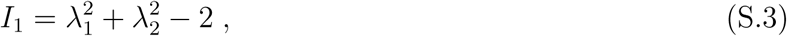

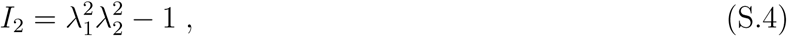

where *κ*_*s*_ and *κ*_*α*_ are the shear and dilation moduli, *λ*_1_, *λ*_2_ are the local principal in-plane stretch ratios (see [51] for calculation procedure), and *W*^*S*^ = ∫*dA w*^*S*^. The shape of the discocyte membrane is approximated by a number *N*_*f*_ of flat triangular faces, and *W*^*S*^ is numerically calculated based on a finite element method (FEM) approach as

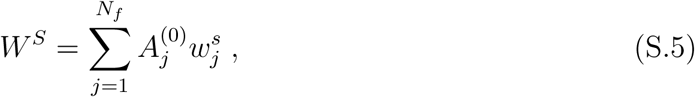

where 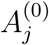 is the undeformed area of face *j*. The bending energy of the RBC membrane is numerically calculated as

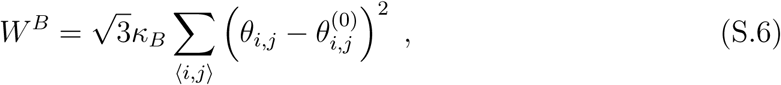

where *κ*_*B*_ is the bending modulus, *θ*_*i,j*_ is the angle between the normals of two neighbouring faces *i* and *j*, and 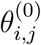 is the same angle for the undeformed membrane. Finally, we penalise deviations of the total membrane surface area and volume by defining two additional energy contributions:

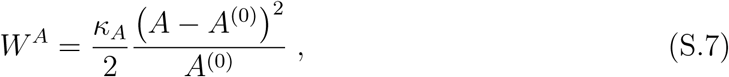

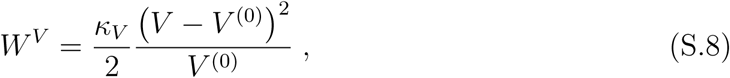

where *κ*_*A*_, *κ*_*V*_ are the surface area and volume moduli, *A* and *A*^(0)^ are the current and undeformed membrane surface areas, and similarly with *V*. The principle of virtual work yields the force acting on each membrane vertex *i* at position 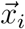 through

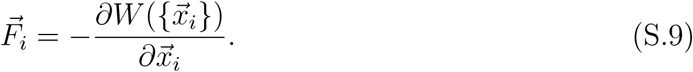

The immersed boundary method [62] is used to couple the fluid and membrane dynamics. The fluid velocity is interpolated at the positions 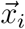 of the RBC mesh vertices, and a forward-Euler scheme is used to advect the vertices to satisfy the no-slip condition. The vertex forces 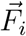 are spread to the lattice where they are used as input to the forcing term in the LBM, which ensures local momentum exchange between the membrane and the fluid. See [29] for a detailed numerical analysis of the algorithm.

The RBC model contains five parameters (*κ*_*s*_, *κ*_*α*_, *κ*_*B*_, *κ*_*A*_, and *κ*_*V*_). While *κ*_*s*_ and *κ*_*B*_ are known from experiments (see review in [63]), the exact values of the three remaining parameters are chosen to ensure that local area, total surface area and volume drift are constrained within a few percent and simulations are stable (see analysis is [29, 51]). Table S7 summarises all the parameters in the model.

### A.3 CFL width calculation

To calculate the CFL width in channel 1 of the domains in Figure 3, let us consider a vessel cross-section of diameter *d* at distance *l* downstream from the first bifurcation in the network. The RBC density, *ϕ*(*r, θ, l, t*), is 1 if there is a RBC at time *t* occupying the point with radial coordinate *r* and angular coordinate *θ* of the cross-section and 0 otherwise. The average RBC density flux Φ(*l*) going through the cross-section is

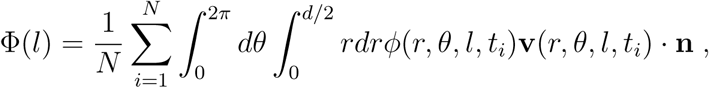

where **v** is the fluid velocity, **n** is the cross-section normal vector and *N* is the number of simulation time steps in the average (0.5 s of real time simulation sampled every 0.0215 s, *N* = 23, in our case).

We define *χ* = 0.01 as the average fraction of RBC density flux crossing the CFL. Now we are able to numerically determine the local CFL width *W* (*l, θ*): consider a 2D-cone centered and contained in the cross-section with orientation *θ* and size Δ*θ* = *π/*2. The width *W* (*l, θ*) is the distance such that

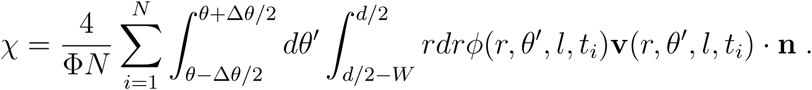

Since we are only interested in the spatial evolution of the CFL, the specific value of *χ* used in the definition is arbitrary. The choice of *χ* will change the width of the CFL after symmetry recovery, but it will not affect the local characterisation of the CFL spatial evolution after a bifurcation. For example, for any value of *χ*, the CFL recovery distance can be calculated as the shortest distance *l* for which the CFL width *W* do not depend on coordinate *θ*.

### A.4 Hybrid model of oxygen transport in vascularised tissue

#### A.4.1 Choice of vessel diameters and branching angles in vascular networks

In the branched networks used, we fix the diameter of the inlet vessel so that *d*_*inlet*_ = 100 *µ*m. The diameters of the two child vessels (*d*_*α*_ and *d*_*β*_) are assumed to be equal and determined from the diameter of the parent vessel (*d*_*P*_) via Murray’s law [42] so that:

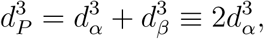

in which case

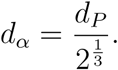

Since the network is symmetric about its central axis, vessels on the converging side have the same diameters as those of the same generation on the diverging side (see Supplementary Figure S4a). For all simulations the networks have 6 generations of vessels. The length *L* of a vessel segment in a given network is related to its diameter *d* via *L* = *λd*, where the positive constant *λ* is network-specific.

For complete specification of the network geometry, in two-dimensional cartesian geometry, it remains to embed the network in a spatial domain. This is achieved by specifying either the branching angles, or (equivalently) the lengths of the projections of the vessels on the *y* axis. Denoting by 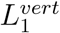 the length of the projection of a vessel of generation 1, the lengths of the projections of vessels of generation *i* > 1 are given by 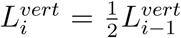. As a result, the vertical size of the domain will not exceed 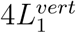 for any number of generations. Finally, we require 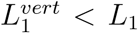 = length of vessels of generation 1. In our simulations, we fix 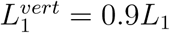 to ensure adequate spatial extent in the *y*-direction.

#### A.4.2 Poiseuille’s law and the Fahraeus-Lindquist effect

We simulate flow in the branched networks by following the approach of Pries *et al.* [54]. For blood vessels of length *L* and diameter *d*, we assume Poiseuille’s law

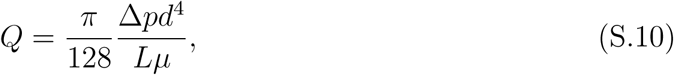

where *Q* is the vessel flow rate, Δ*p* is the pressure drop along the vessel and *µ* is the effective viscosity of blood [53]. Following [64] we assume that the blood viscosity depends on vessel diameter and haematocrit via the empirical relationship:

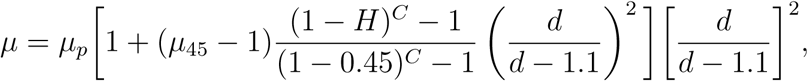

where *µ*_*p*_ is the plasma viscosity, *H* is the vessel discharge haematocrit,

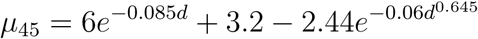

and

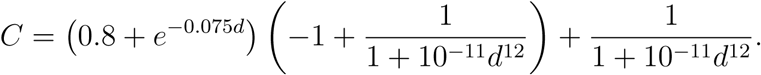

Introducing signed flow rates 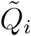 for the sake of brevity, we impose conservation of blood and haematocrit at each network bifurcation, so that

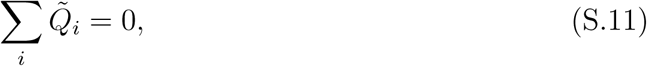

and

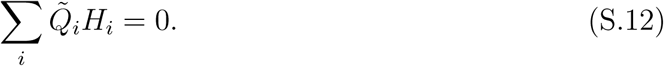

In (S.11) and (S.12) we sum over the three vessels that meet at that bifurcation. At diverging bifurcations, we impose a HS rule: we use (1) from the main text when CFL memory effects are neglected and (2) from the main text when they are included. Denoting by *N*_*B*_ the number of network bifurcations and *N*_*V*_ the number of vessels, we have *N*_*B*_ unknown pressures *P, N*_*V*_ unknown flow rates *Q* and *N*_*V*_ − 1 unknown haematocrit levels (the inlet haematocrit being prescribed) - altogether *N*_*B*_ + 2*N*_*V*_ − 1 unknowns. At the same time, we impose Poiseuille’s law ((S.10)) for every vessel (*N*_*V*_ times), conservation of blood ((S.11)) and haematocrit ((S.12)) at every bifurcation node (*N*_*B*_ times), and an HS rule at all diverging bifurcations (*N*_*B*_*/*2 times), yielding a total of *N*_*V*_ + 5*N*_*B*_*/*2 algebraic equations. Since every bifurcation connects 3 vessels, we have *N*_*V*_ = (3*N*_*B*_ + 2)*/*2, where every vessel is counted twice, except for the inlet and outlet vessels (+2 in the numerator). From this, it follows that the number of equations (*N*_*V*_ + 5*N*_*B*_*/*2) equals the number of unknowns (*N*_*B*_ + 2*N*_*V*_ − 1).

#### A.4.3 Oxygen distribution in tissue

In this section, we determine the oxygen concentration *c* in the tissue. Following [13], we assume that the dominant processes governing its distribution are delivery from the vessel network (via one-way coupling with (S.12) and the haematocrit models (1) or (2), *i.e. c* depends on *H*_*l*_ but not vice versa), diffusive transport through the tissue, and consumption by cells in the tissue. We focus on the long time behaviour and, therefore, adopt a quasi-steady state approximation [65]

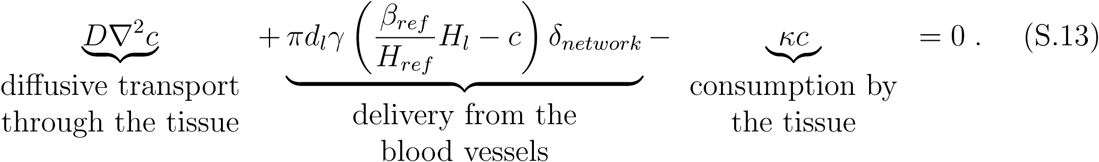

In (S.13), the positive constants *D, γ* and *κ* represent the diffusion coefficient for oxygen in the tissue, the vessel permeability to oxygen, and the rate at which it is consumed by cells in the tissue. The vessel network is represented by a collection of Dirac point sources *δ*_*network*_ where

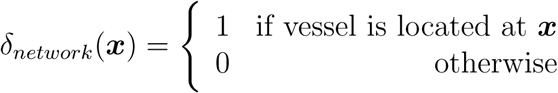

and for any ***x*** satisfying *δ*_*network*_(***x***) = 1, *d*_*l*_ and *H*_*l*_ are the diameter and haematocrit of the vessel at that location (where the latter has been calculated as described in the previous section). The constant *β*_*ref*_ represents the oxygen concentration of a reference vessel containing haematocrit *H*_*ref*_ (here we fix *H*_*ref*_ = 0.45, the inlet haematocrit) and we suppose that the oxygen concentration of a vessel with haematocrit *H*_*l*_ is *β*_*ref*_ *H*_*l*_*/H*_*ref*_. In (S.13) we assume that the oxygen is supplied by vessels to the tissue at a rate which is proportional to their circumference *πd*_*l*_, the vessel permeability *γ*, and *β*_*ref*_ *H*_*l*_*/H*_*ref*_ − *c*. Finally, we have *β*_*ref*_ = *c*_*stp*_*p*_*ref*_ *α*_*eff*_, where *c*_*stp*_ denotes an ideal gas concentration at standard temperature and pressure, *p*_*ref*_ denotes the reference partial pressure at the inlet vessel, and *α*_*eff*_ denotes the volumetric oxygen solubility [66]. A summary of the parameter values used to solve (S.13) is presented in Supplementary Table S8.

### A.5 Derivation of, and justification for, the HS model with CFL memory

#### A.5.1 Parameter dependencies in HS model without memory from [39]

The dependencies of the parameters *A, B* and *X*_0_ (see (1) from the main text) on the diameters of the parent and child vessels (*d*_*P*_, *d*_*α*_ and *d*_*β*_, respectively), and the discharge haematocrit *H*_*P*_ in the parent vessel were first introduced in [11, 54] and later adjusted in [39] to achieve a better approximation under extreme combinations of *d*_*α*_, *d*_*β*_, *d*_*P*_ and *H*_*P*_. We will use the functional forms from [39], which read

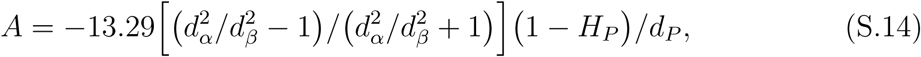

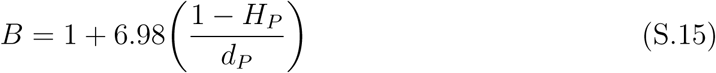

and

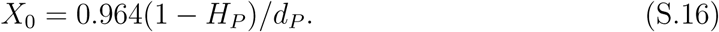

These functional forms assume that *d*_*P*_ is dimensionless and given by 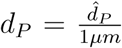, where 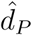 is the dimensional diameter. We maintain this convention throughout this section.

#### A.5.2 HS model with memory

##### Simplifying assumptions

Before we explain how we extend the model from [39] to incorporate memory effects, we comment on its main simplifying assumptions. At present, our model does not include any information on local flow rate. Furthermore, the current model does not account for the angle defined by the planes containing the current and previous bifurcation in the network. These simplifying assumptions could easily be relaxed.

##### Rewriting of the model

In this section, we rewrite the HS model with memory effects ((2) from the main text) in terms of discharge haematocrit levels *H* and flow rates *Q* experienced by the vessels belonging to a given bifurcation ((3) from the main text). The definitions of *FQ*_*E,f*_ and *FQ*_*B,f*_ can be written as:

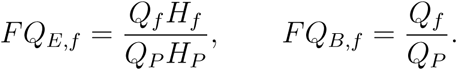

Substituting these expressions into (2) from the main text gives:

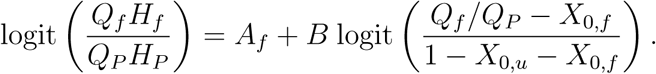

Recalling that logit(*x*) = ln (*x/*(1 − *x*)), we have

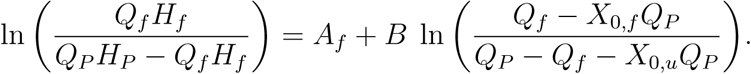

Appealing to conservation of blood (overall)

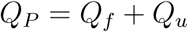

and RBCs (in particular)

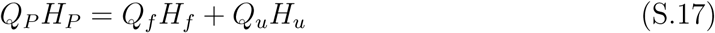

at diverging bifurcations, we arrive at

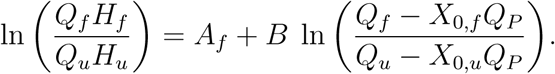

This equation can also be written as

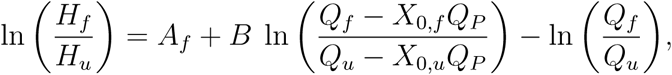

which yields

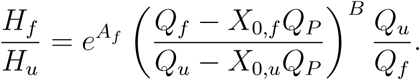

##### Choice of parameter values and CFL recovery function

Here we introduce the functional forms for *A*_*f*_, *X*_0,*f*_ and *X*_0,*u*_, using empirical data to justify our choices. Guided by the dependence of *A* on the network branching history described in [11] (see Figure 7 therein), we propose

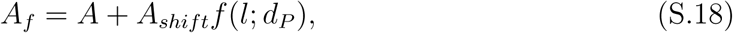

where *A* is given in (S.14), the positive constant *A*_*shift*_ corresponds to the maximum CFL disruption effect, and the function *f* (*l*; *d*_*P*_) describes how the recovery of the CFL depends on the distance *l* to the previous bifurcation and the diameter *d*_*P*_ of the parent vessel^3^.

For parameter *A*, we only have access to the scattered data with respect to the regressor from [11, 54] (as opposed to the regressor from [39]), which reads

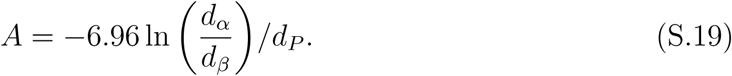

Using the extreme values of *A* in these data (see Supplementary Figure S8c), we estimate *A*_*shift*_ = 0.5. Note that in branching networks with every pair of child vessels having equal radii, both [54] and [39] yield *A* = 0. Thus, for our networks, the choice of *A* does not affect *A*_*f*_ at all (see (S.18)).

For simplicity, we model the CFL recovery using an exponential function

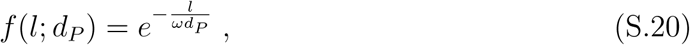

where *ω* controls the temporal dynamics of CFL recovery. From [37], we note that the CFL width is (approximately) 90% recovered at a distance *l*_90_ = 10*d*_*P*_ from the previous bifurcation (see also Figure 3g). Accordingly, we choose *ω* so that

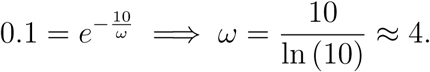

Guided by the dependence of *X*_0,*f*_ on flow history described in [11], we propose

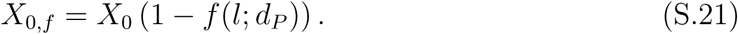

Assuming, as a first approximation, that *X*_0,*f*_ + *X*_0,*u*_ is constant and independent of the distance to the previous bifurcation (see Figure 3g), we define

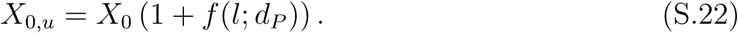

#### A.5.3 Validation of the HS model with memory

We validate the HS model with memory by comparing its predictions with results from the RBC simulations in the double-t geometry. We assume that all vessels have the same diameter (*d* = 33 *µ*m), and that the flow rate splits evenly at both bifurcations. If we assume further that the CFL is fully established at the network inlet vessel, *H*_*inlet*_ = 20%, then (1) from the main text supplies *H*_1_ = *H*_2_ = *H*_*inlet*_ = 20%. We use conservation of RBCs (S.17) and the new HS model (Eq. (3) from the main text) to estimate haematocrit values in the unfavourable and favourable child branches after the second bifurcation (channels 3 and 4, respectively) for varying inter-bifurcation distances *δ*. The results are summarised in Supplementary Table S9. For *δ* = 4*d*, the new HS model predicts haematocrits within 5% relative error of the values calculated from RBC simulations (Table 2). Given the uncertainty in determining discharge haematocrit in the RBC simulations and given that the new model neglects effects due to asymmetric streamline splitting [38], we conclude that our new model provides a good, leading-order approximation to the effects of CFL disruption on HS.

Finally, we compare the CFL evolution dynamics calculated from the RBC simulations (for *θ* = 0 and *θ* = *π*) with those predicted from the proposed evolution of *X*_0,*f*_ and *X*_0,*u*_ ((S.21) and (S.22)). In the absence of a known functional form relating the CFL width *W* and the minimum flow fraction *X*_0_, we define

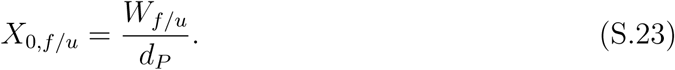

(S.23) is based on the diagram in Supplementary Figure S8a and the assumptions of a cross-sectionally uniform velocity profile within a one-dimensional vessel cross-section. Combining (S.16), (S.23), (S.21) and (S.22), we conclude

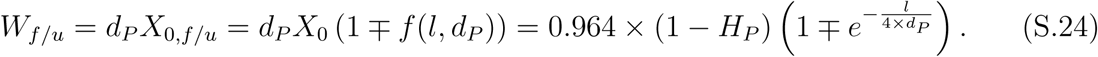

We remark that for a well-established CFL (i.e. *l* → *∞*), (S.24) predicts (noting that channel 1 serves as the parent vessel for the second bifurcation and estimating *H*_*P*_ = 0.2 from Supplementary Table S6) a CFL width of about 0.77 *µ*m, whereas our RBC simulation predicts a value of approximately 1.8 *µ*m (see Supplementary Figure S8b). We postulate that this discrepancy is caused by our oversimplification of the relationship between the CFL width and the minimum flow fraction ((S.23)). Nevertheless, we can adjust (S.24) so that it is consistent with the established CFL width of 1.8 *µ*m by writing

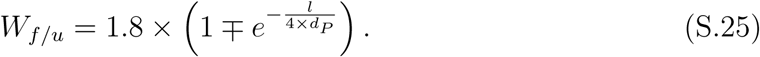

In this case the CFL evolution (for *θ* = 0 and *θ* = *π*) follows a trend similar to that observed in our RBC simulations (Supplementary Figure S8b). In particular, our assumption that *l*_90_ = 10*d*_*P*_ is in good agreement with our simulation results (see dashed line in Supplementary Figure S8b).

### A.6 Explanation of higher mean oxygen values for small *λ*

We observed that CFL disruption effects increase the mean oxygen concentration in the chosen network (Supplementary Figure S6). Here, we provide an explanation of this phenomenon.

We define

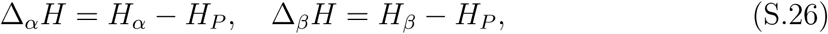

where *P* is the parent branch and *α* and *β* are the child branches of any diverging bifurcation. Conservation of blood and RBCs at this bifurcation then yields

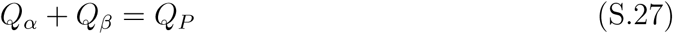

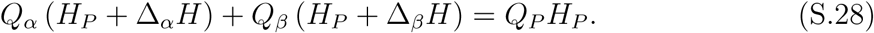

Combining (S.28) and (S.27) supplies

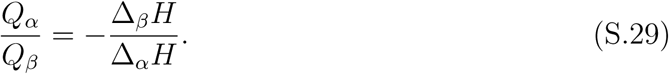

We deduce that, at diverging bifurcations, the haematocrit level in the child branch with higher flow rate deviates less (in absolute value) from the haematocrit in the parent vessel than the branch with lower flow rate.

We note further that all paths connecting the inlet and outlet vessels in the direction of blood flow in a given network are topologically and geometrically equivalent. Therefore, heterogeneity in haematocrit splitting arises solely from CFL disruption effects. If haematocrit is elevated in one of the child branches, its impedance will increase, and, as a result, it will receive a lower flow rate.

Combining these two effects, we see that, in the chosen networks, haemoconcentration in any child branch is more significant than haemodilution in its sibling. As a consequence, and given that the strength of the oxygen source term in (S.13) is a linear function of *H*, we observe higher mean oxygen levels when the effects of CFL disruption are taken into account (especially for small *λ*). Future work will investigate this effect by making source term a function of RBC mass flux (*i.e. QH*) or relaxing the assumption that the RBCs have infinite oxygen carrying capacity.

## B Supplementary tables/figures

Supplementary figures and tables supporting the main manuscript text and supplementary information:

- Supplementary Tables S1–S9.
- Supplementary Figures S1–S8

**Table S1:**
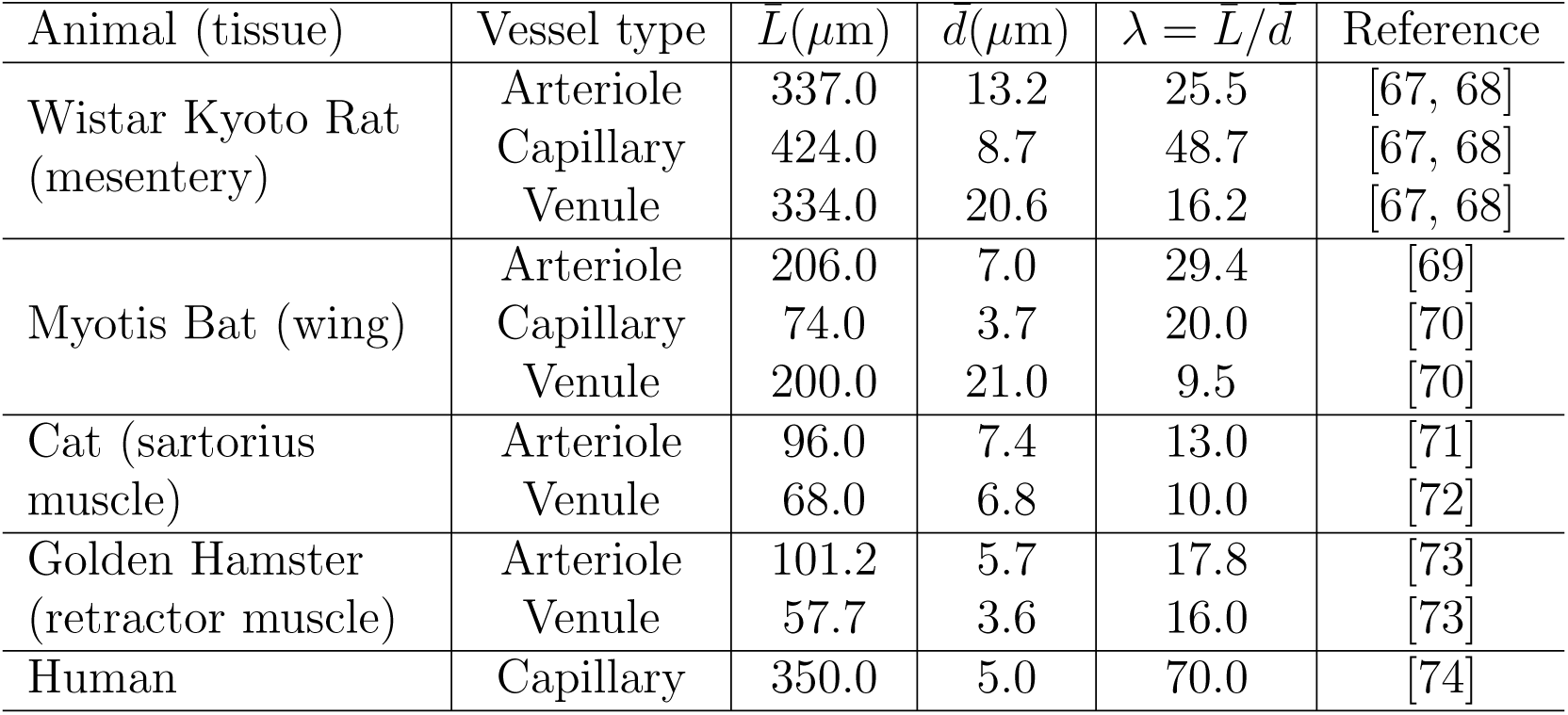
Average vessel lengths and diameters reported in a variety of tissues under physiological conditions.

| Animal (tissue) | Vessel type | $\bar{L}(\mu\text{m})$ | $d(\mu\text{m})$ | $\lambda = \bar{L}/d$ | Reference |
| --- | --- | --- | --- | --- | --- |
| Wistar Kyoto Rat<br>(mesentery) | Arteriole | 337.0 | 13.2 | 25.5 | [67, 68] |
|  | Capillary | 424.0 | 8.7 | 48.7 | [67, 68] |
|  | Venule | 334.0 | 20.6 | 16.2 | [67, 68] |
| Myotis Bat (wing) | Arteriole | 206.0 | 7.0 | 29.4 | [69] |
|  | Capillary | 74.0 | 3.7 | 20.0 | [70] |
|  | Venule | 200.0 | 21.0 | 9.5 | [70] |
| Cat (sartorius<br>muscle) | Arteriole | 96.0 | 7.4 | 13.0 | [71] |
|  | Venule | 68.0 | 6.8 | 10.0 | [72] |
| Golden Hamster<br>(retractor muscle) | Arteriole | 101.2 | 5.7 | 17.8 | [73] |
|  | Venule | 57.7 | 3.6 | 16.0 | [73] |
| Human | Capillary | 350.0 | 5.0 | 70.0 | [74] |

**Figure S1:**
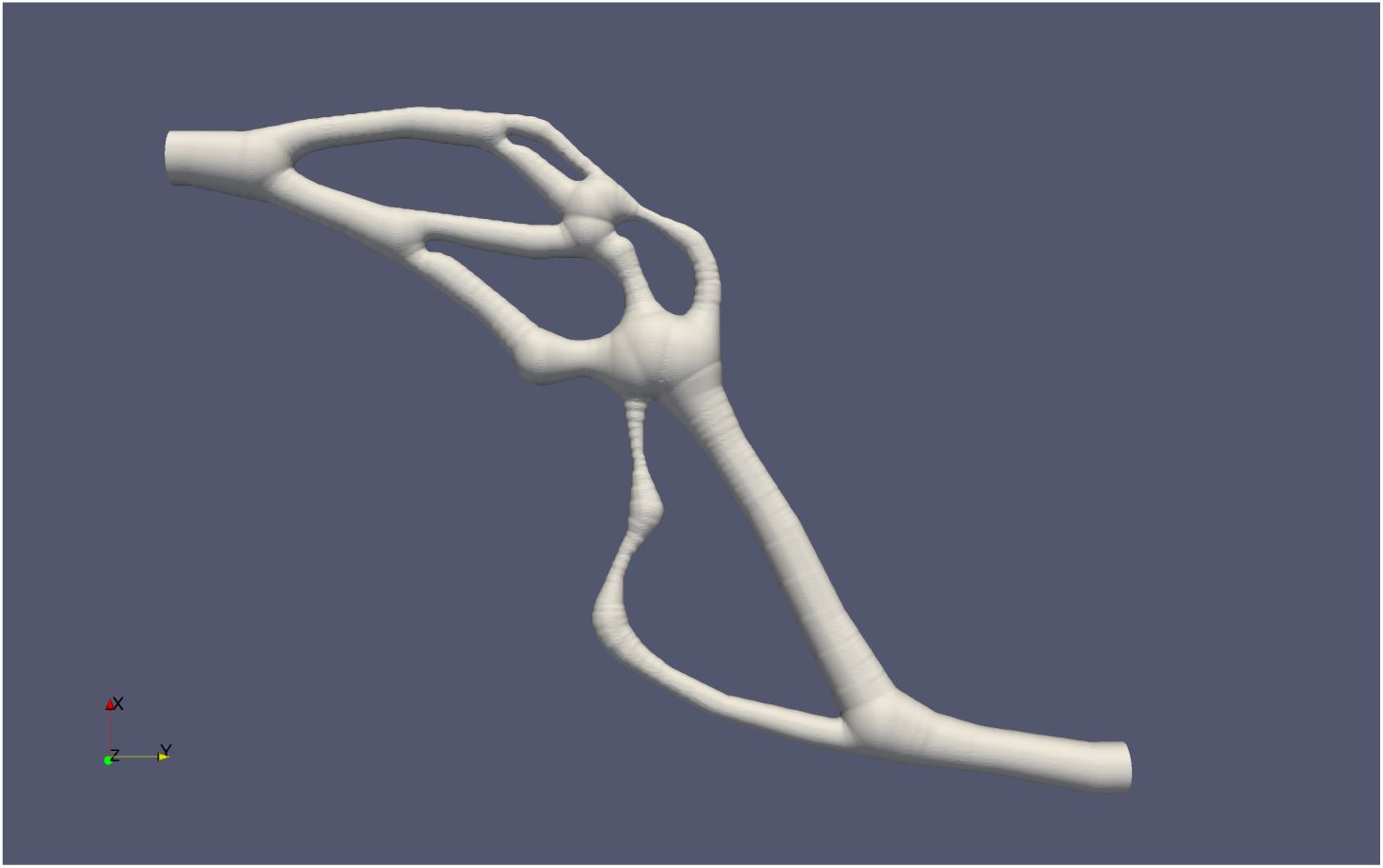
Realistic capillary network reconstructed from MC38 tumour vascular network.

**Figure S2:**
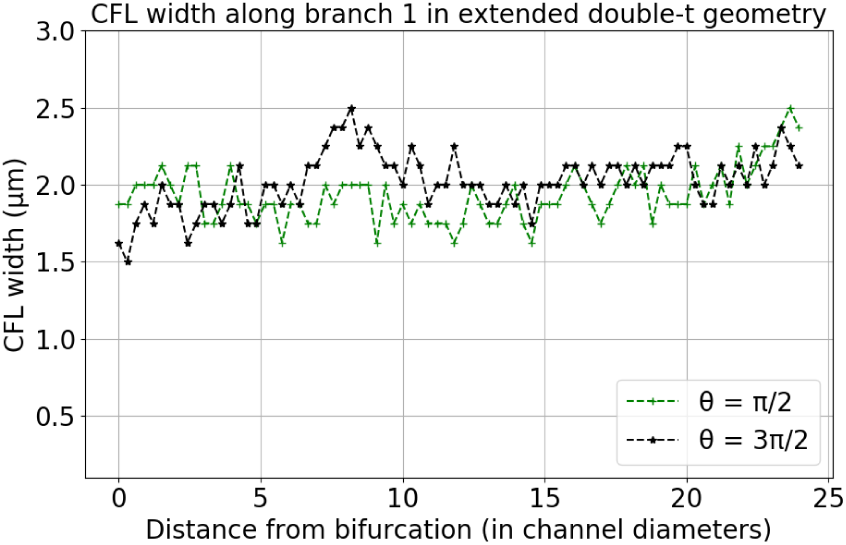
CFL channel 1 double-t geometry perpendicular to bifurcation planes.

**Table S2:**
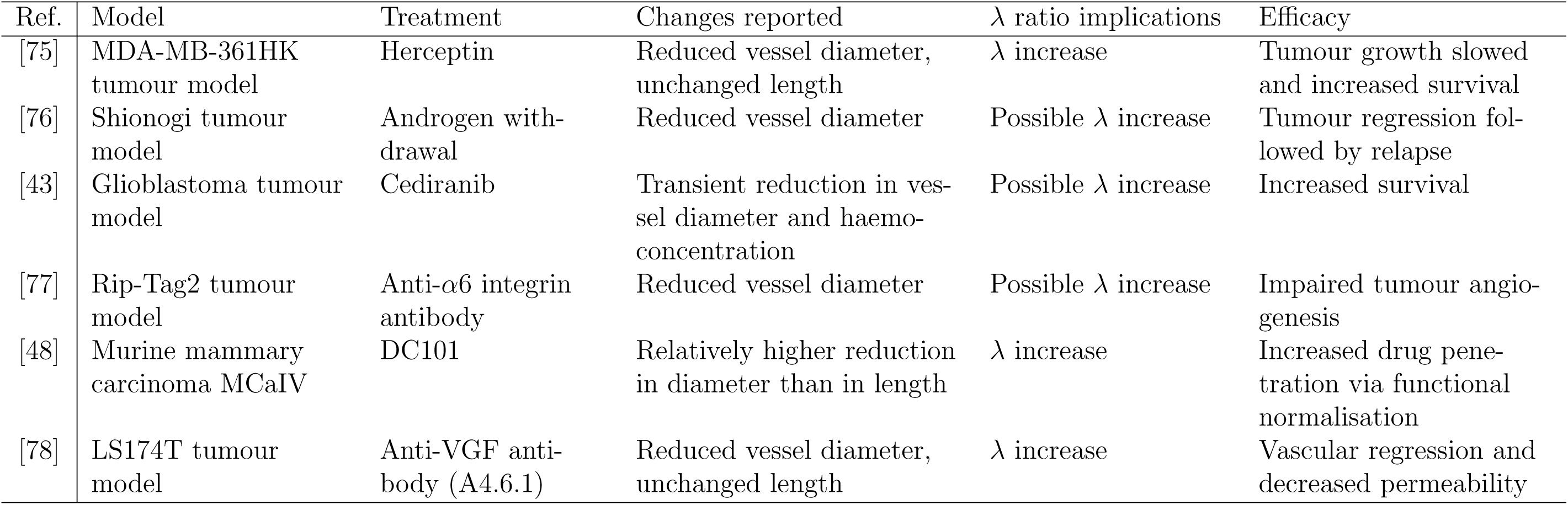
Previous studies reporting morphological or haemodynamic changes following vascular normalisation therapy.

**Figure S3:**
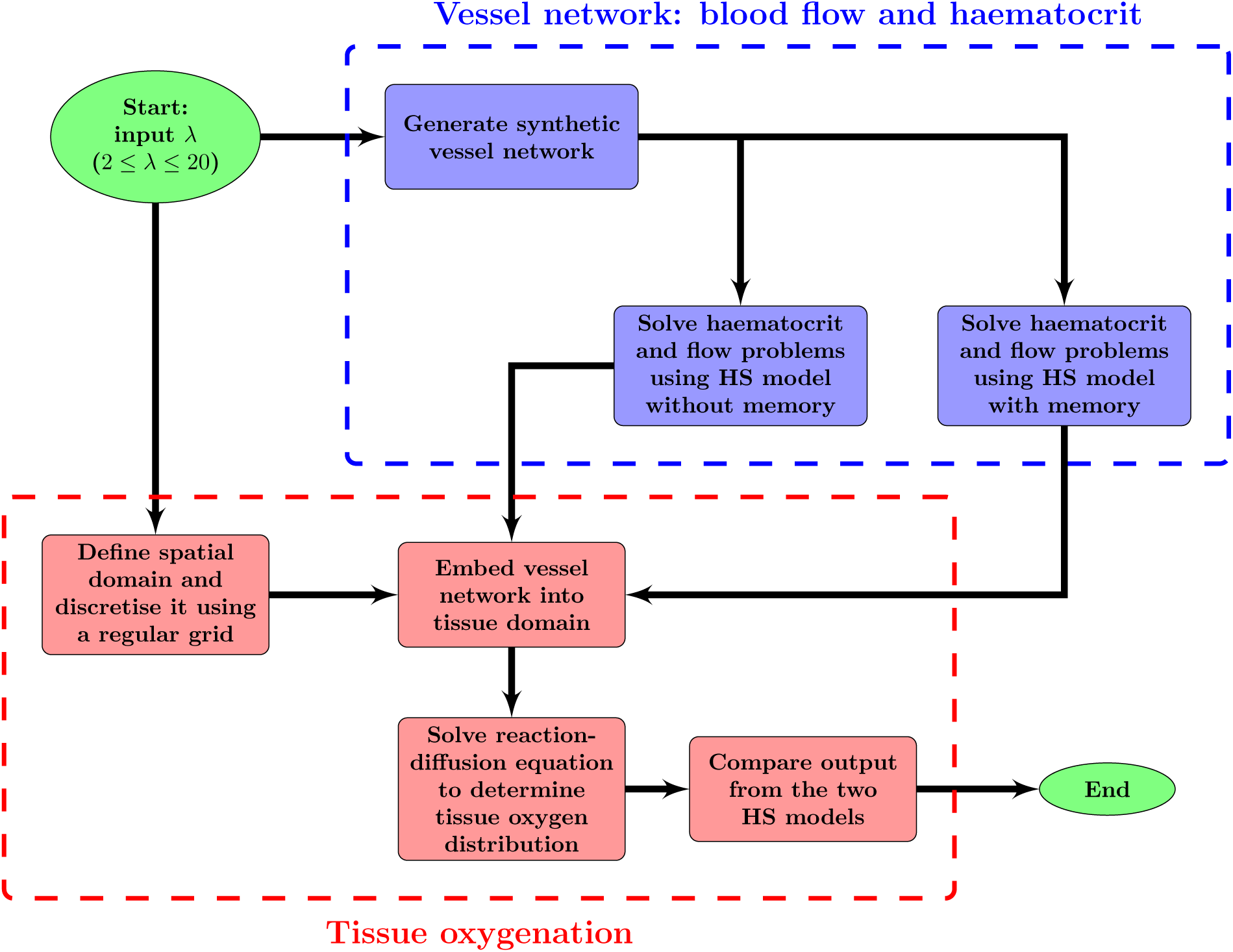
Flow chart summarising the main components of our hybrid model for tissue oxygen perfusion, as implemented within Microvessel Chaste (see [13]).

**Table S3:**
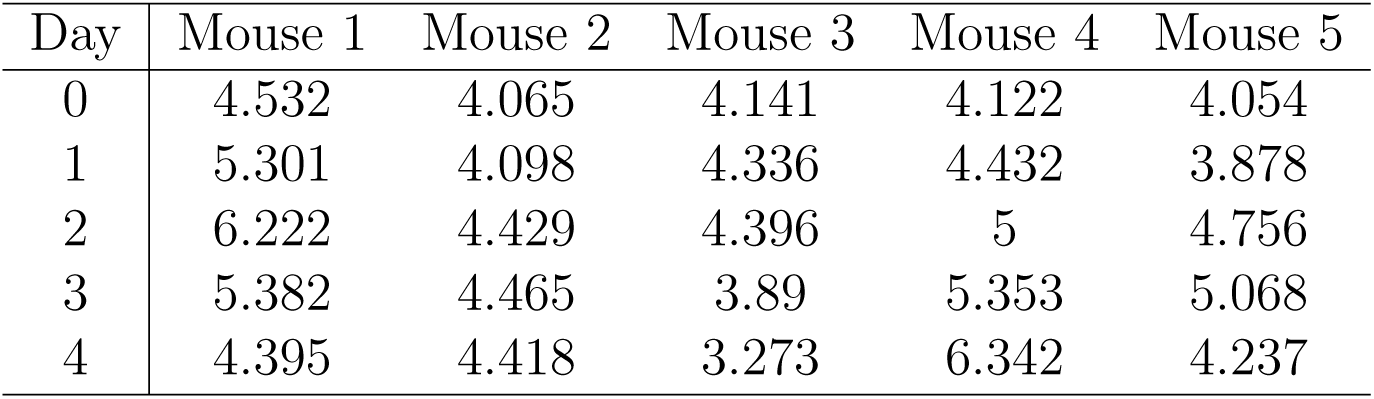
*λ* values measured in MC38 tumours following DC101 treatment over time.

**Figure S4:**
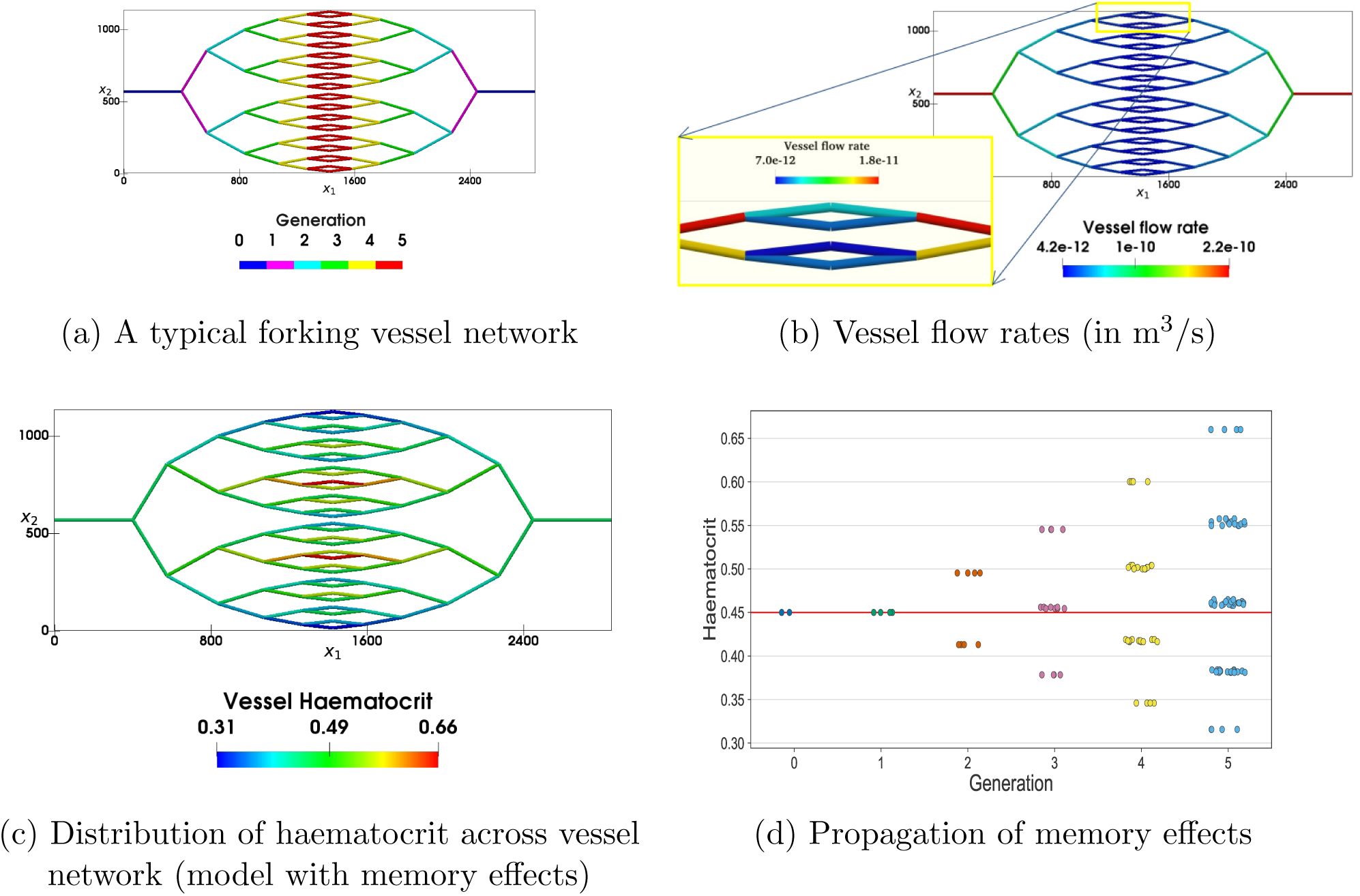
(a) A typical symmetric forking network with 6 generations of vessels. (b) Flow rates almost halve between consecutive vessel generations. However, small differences in flow rates between child vessels arise due to non-uniform haematocrit splitting (HS), as can be observed in the inset (note that the range of the colour bar has been adjusted to represent only the selected vessels). (c) Differences in the predicted haematocrit levels of child vessels (within a single vessel generation) become more pronounced as the generation number increases. (d) For the new HS model, the haematocrit distribution becomes more disperse as the number of bifurcations included in the network increases (the red horizontal line represents the predicted haematocrit when memory effects are neglected and haematocrit is distributed uniformly across the network). Each circle corresponds to a single vessel and different colours correspond to different vessel generations.

**Figure S5:**
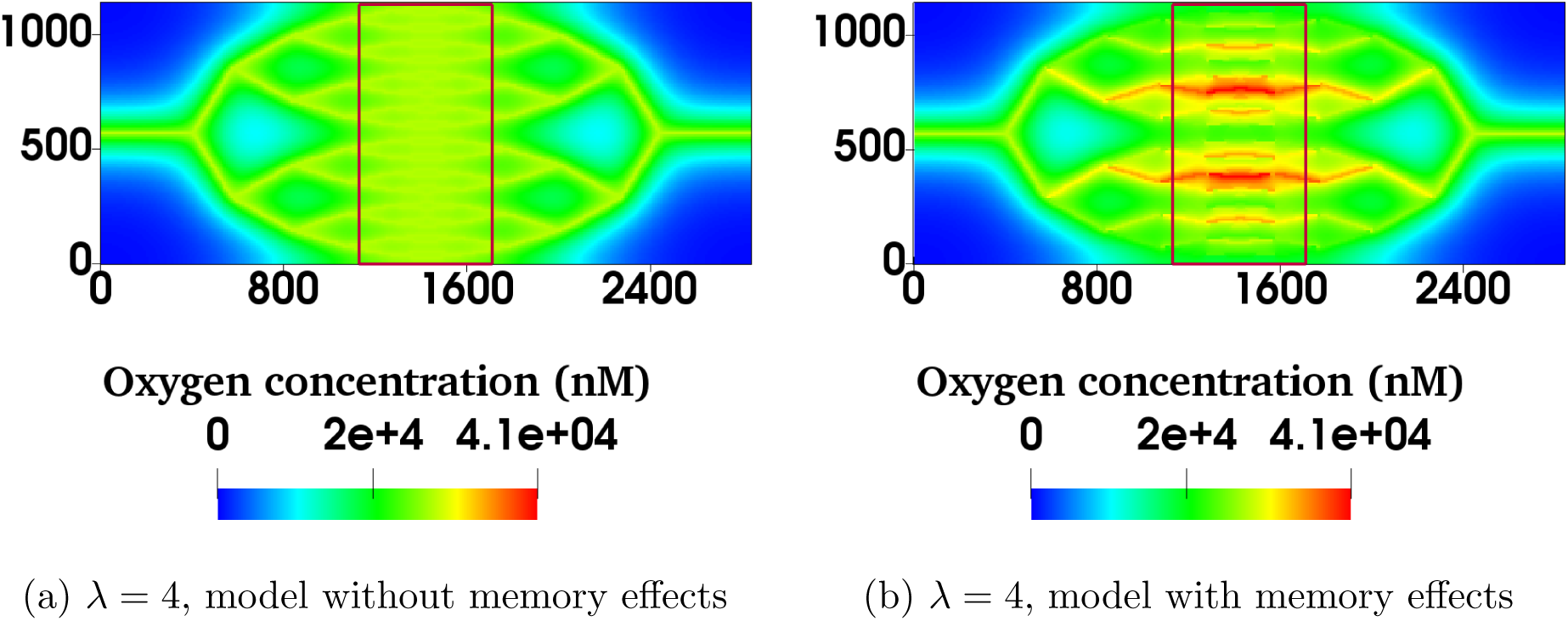
For *λ* = 4.0, the model with memory effects yields more pronounced oxygen heterogeneity (*i.e.* more dispersed oxygen distribution) in the region of interest bounded by red rectangles in (a) and (b) (note that the spatial scales are in microns).

**Table S4:** Timecourse of Pearson’s r-values calculated for different mice at different days of measurement implanted with the MC38 cell line. Day 0 corresponds to the day of the first measurement, when the tumour reached a specified size (4mm in diameter; see the main text). The corresponding values of 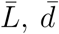 and *λ* are reported in the main text (see Table 1). The missing datum for tumour 3 on Day 3 is due to the laser on the microscope failing during imaging.

| Day | 1 | 2 | 3 | 4 | 5 | 6 |
| --- | --- | --- | --- | --- | --- | --- |
| 0 | 0.05 | -0.06 | -0.07 | 0.10 | -0.13 | 0.00 |
| 1 | 0.03 | -0.05 | -0.07 | -0.07 | -0.13 | -0.00 |
| 2 | -0.08 | -0.07 | -0.06 | -0.17 | -0.19 | 0.02 |
| 3 | -0.09 | -0.09 | - | -0.14 | -0.13 | -0.02 |
| 4 | -0.14 | -0.11 | -0.08 | -0.17 | -0.12 | -0.07 |
| 5 | - | -0.09 | - | - | -0.07 | -0.11 |
| 6 | - | -0.04 | - | - | - | - |
| 7 | - | -0.08 | - | - | - | - |

**Table S5:**
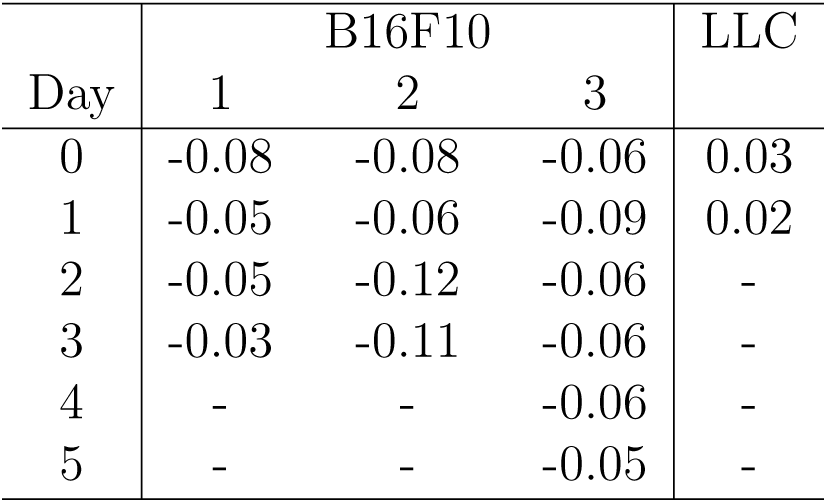
Timecourse of Pearson’s r-values for mice implanted with the B16F10 and LLC cell lines. Day 0 corresponds to the day of the first measurement, when the tumour reached a specified size (4mm in diameter; see the main text). The corresponding values of 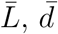and *λ* are reported in the main text (see Table 1).

**Figure S6:**
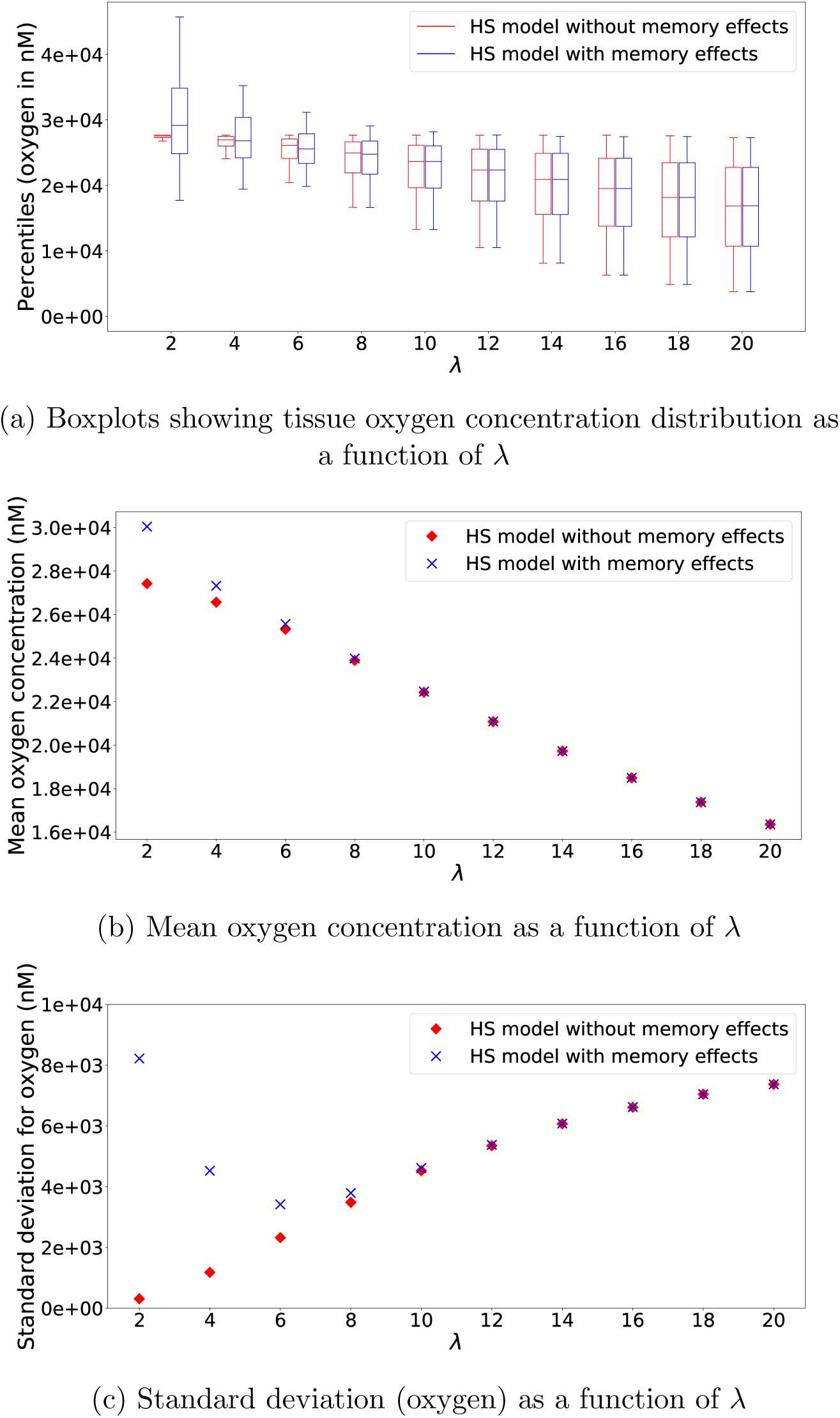
Summary statistics illustrating how for a vessel network with 6 generations its *λ* value and the HS model affect tissue oxygenation. (a) Boxplots showing how the tissue oxygen distribution changes as *λ* varies for the two different HS rules. (b) Mean oxygen concentration increases as *λ* decreases (and the vessel density increases). (c) Standard deviation in the tissue oxygen concentration increases with *λ* when memory effects are neglected ((1) from the main text). When memory effects are considered ((2) from the main text), the standard deviation increases for small *λ* values. The mean and standard deviation for the two models converge for large *λ* values. Model parameter values as per Supplementary Table S8.

**Figure S7:**
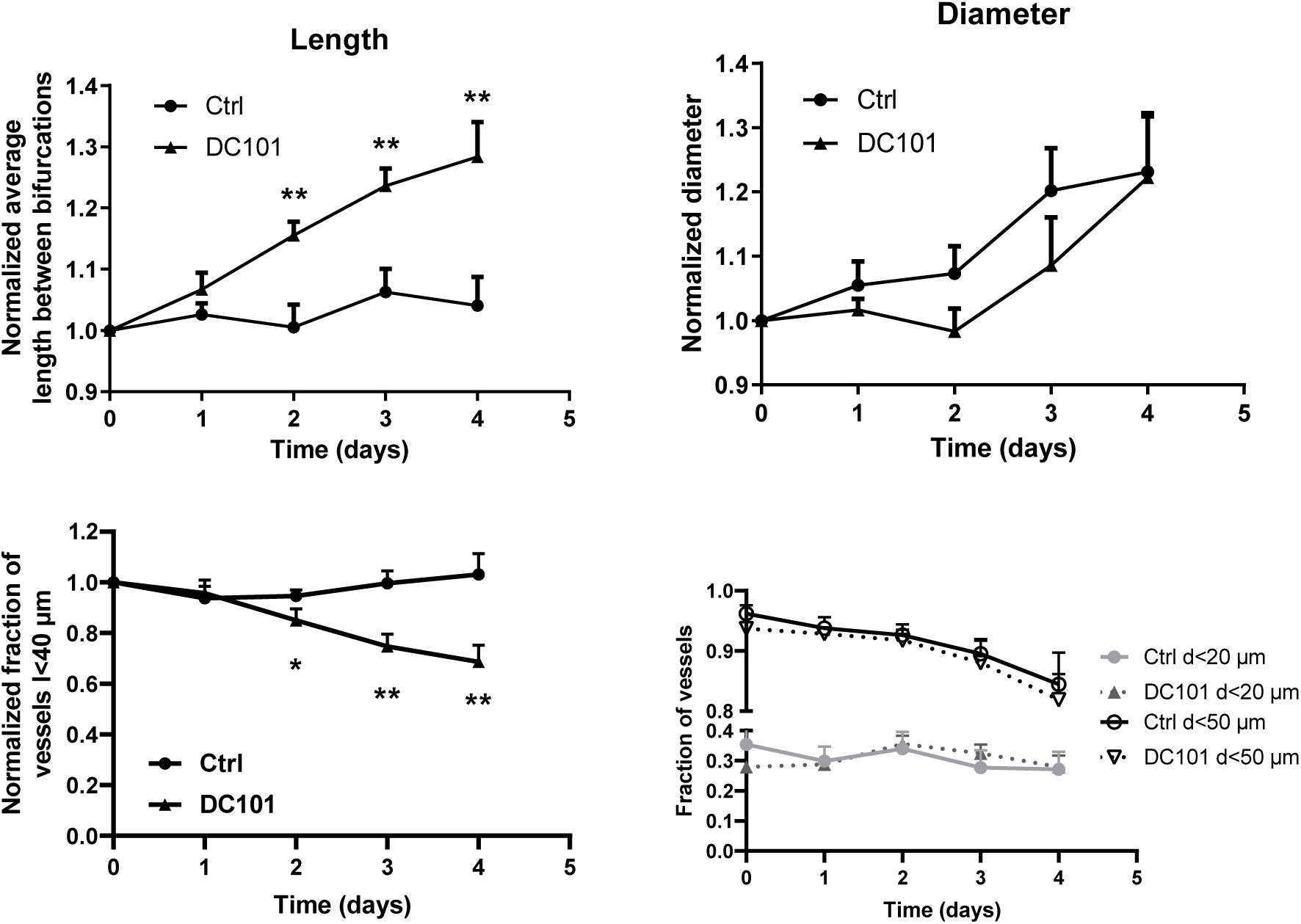
Vascular phenotypes in MC38 tumours over time following DC101 treatment compared with control (n=5). * *p* < 0.05, ** *p* < 0.01.

**Table S6:** Parameters in RBC simulations in synthetic capillary networks

| Parameter | Description | Value | Reference |
| --- | --- | --- | --- |
| $d$ | Cylindrical channel diameter | $33 \mu\text{m}$ | Current study |
| $L'$ | Inlet/outlet channel length | $25d$ | [37] |
| $\delta$ | Distance between branching points | $4d, 25d$ | [37], current study |
| $\bar{v}_{inlet}$ | Inlet mean velocity | $600 \mu\text{m/s}$ | [18] |
| $H_{inlet}$ | Inlet discharge haematocrit | 20% | [18] |

**Table S7:**
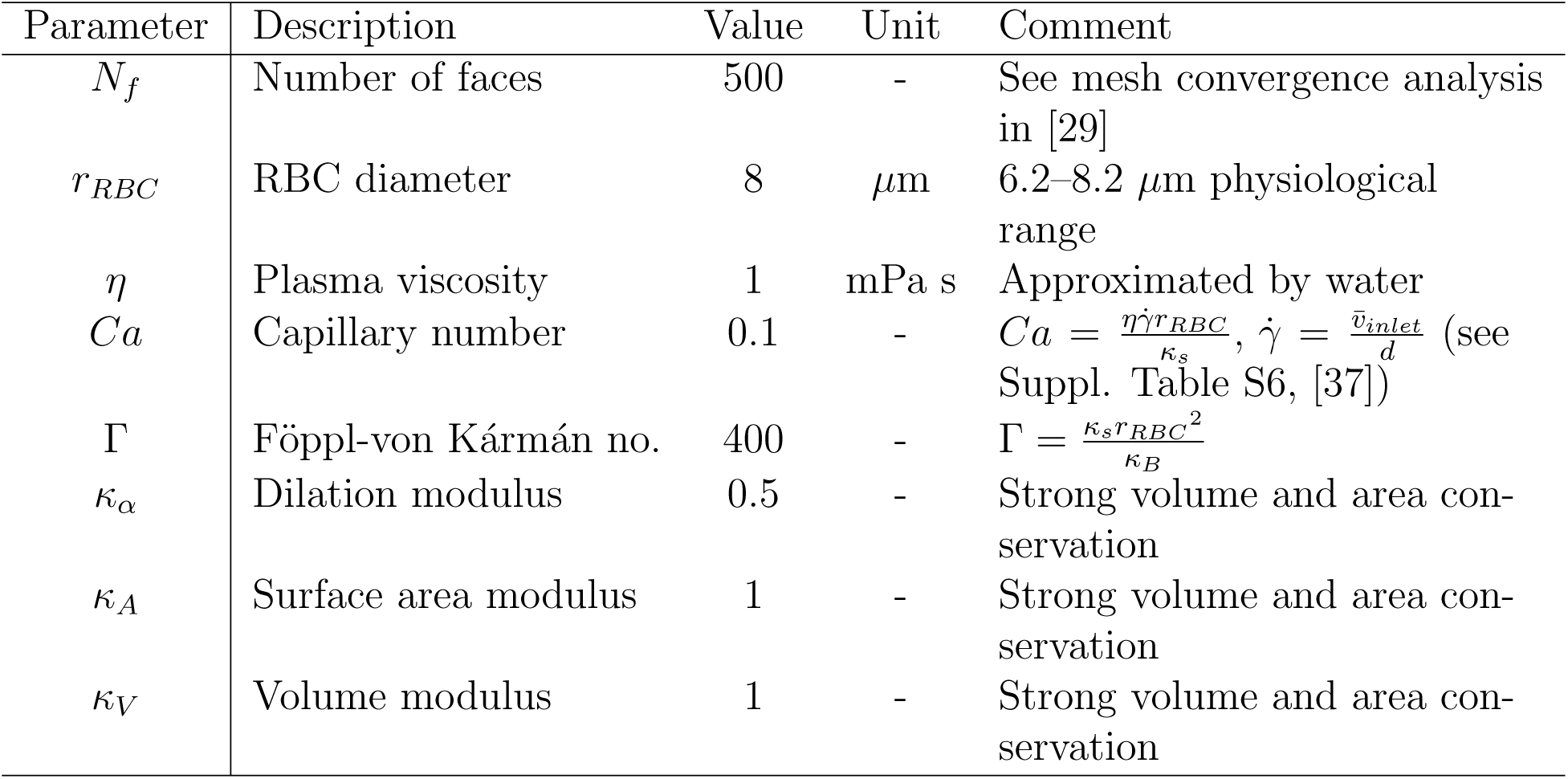
Parameters used in RBC simulation algorithm. The number of faces is chosen in such a way that the average edge length of a triangular element matches the grid spacing of the fluid lattice. The value of the capillary number is representative of typical flows in the microcirculation. The adopted value of the Föppl-von Kárman number matches the intrinsic property of healthy RBC membranes. The remaining moduli are chosen in such a way that the local area, total surface area and volume of the RBCs are constrained within a few percent while simulations remain numerically stable.

**Table S8:** Parameters used to simulate tissue oxygen.

| Parameter | Description | Value | Unit | Reference |
| --- | --- | --- | --- | --- |
| $D$ | Diffusivity | 0.00145 | $\text{cm}^2 \text{min}^{-1}$ | [79] |
| $\kappa$ | Consumption rate | 13.0 | $\text{min}^{-1}$ | [79] |
| $\gamma$ | Vessel permeability | 6.0 | $\text{cm min}^{-1}$ | [79] |
| $c_{stp}$ | Ideal gas concentration | $\frac{1}{0.0224}$ | $\text{mol m}^{-3}$ | [80] |
| $p_{ref}$ | Reference partial pressure | 20 | mmHg | [79] |
| $\alpha_{eff}$ | Volumetric solubility | $3.1 \times 10^{-5}$ | $\text{mmHg}^{-1}$ | [66] |
| $H_{inlet}$ | Inlet haematocrit | 0.45 | - | [81] |
| $d_{inlet}$ | Diameter of inlet vessel | 100 | $\mu\text{m}$ | Estimated from [53] |
| $A_{shift}$ | Maximum CFL disruption effect | 0.5 | - | Estimated here |
| $\omega$ | Temporal dynamics of CFL recovery | 4 | - | Estimated here |
| $\mu_p$ | Plasma viscosity | $10^{-3}$ | Poiseuille | Similar to [79] |
| $p_{in}$ | Inlet pressure | $3.32 \times 10^3$ | Pa | Similar to [79] |
| $p_{out}$ | Outlet pressure | $2.09 \times 10^3$ | Pa | Similar to [79] |

**Figure S8:**
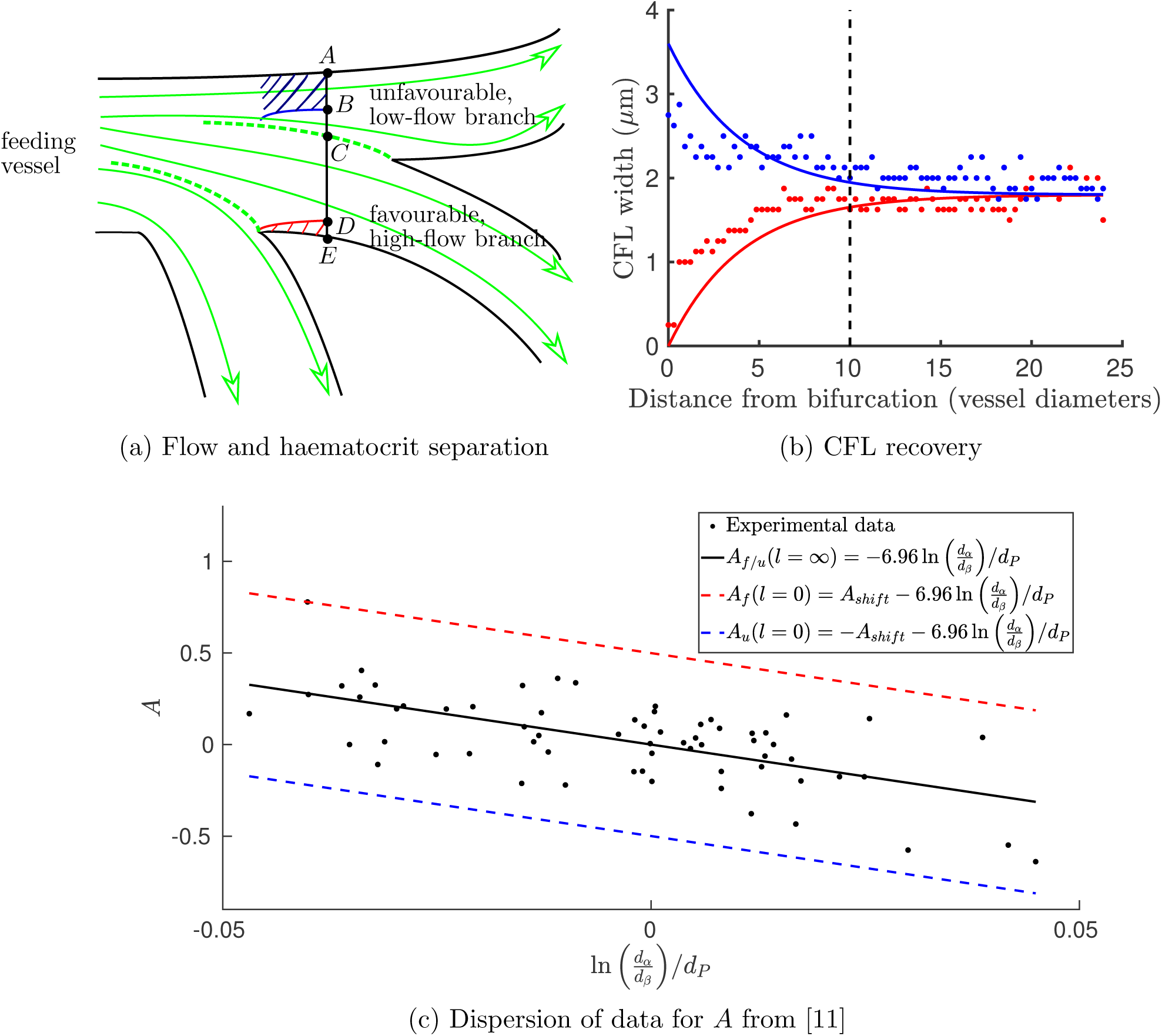
(a) A schematic diagram presenting the geometric intuition behind (blood) flow and haematocrit separation illustrate why two distinct minimum-flow fractions are needed to characterise the favourable and the unfavourable 2 Synthetic vascular networks-branches: ∥*AB*∥ = *X*_0,*u*_, ∥*DE*∥ = *X*_0,*f*_, ∥*AC*∥ = *FQ*_*B,u*_ and ∥*CE*∥ = *FQ*_*B,f*_. Blood flow separation at the two consecutive bifurcations is shown in dotted green, streamlines are sketched with yellow curved arrows, and the CFL recovery on the favourable (unfavourable) side of the parent vessel after the first bifurcation is sketched in red (blue). Whenever *FQ*_*B,f*_ < *X*_0,*f*_ (*FQ*_*B,u*_ < *X*_0,*u*_), the favourable (unfavourable) branch only draws blood from the CFL and it thus receives pure plasma. (b) Model of CFL recovery as described by (S.25) shows similar trends to and is in satisfactory agreement with the CFL width data from RBC simulations in Figure 3g (given the simplifying assumptions). The established CFL width of 1.8 *µ*m chosen by inspection for this particular dataset. (c) Dispersion of values for *A* (reproduced using Figure 6 from [11]) is used with the regression from [54] to estimate the value of *A*_*shift*_ *≈* 0.5 in (S.18), based on deviation from the regression. We assume the CFL disruption to be the primarily cause of this deviation, and thus its maximum (absolute) value should correspond to *l* = 0 in (S.20) (i.e. *f* = 1 in (S.18)).

**Table S9:** Haematocrits predicted by the model with CFL memory effects

| Distance | $H_{inlet}$ | $H_u$ | $H_f$ |
| --- | --- | --- | --- |
| $\delta = 4d$ | 20.0 | 17.7 | 22.3 |
| $\delta = 11d$ | 20.0 | 19.6 | 20.4 |
| $\delta = 18d$ | 20.0 | 19.9 | 20.1 |
| $\delta = 25d$ | 20.0 | 20.0 | 20.0 |

## Footnotes

1 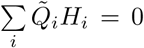, where we sum over all vessels *i* meeting at a given node with haematocrit *H*_*i*_ and signed flow rates 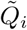 (of magnitude *Q*_*i*_).

2 The dependences of *A, B* and *X*_0_ on the diameters of the participating vessels and on the parent vessel haematocrit are described in Supplementary Information, see (S.14)–(S.16).

3 Consistency of the model requires that *A*_*u*_ = *A* − *A*_*shift*_*f* (*l*; *d*_*P*_).

